# EffectorO: motif-independent prediction of effectors in oomycete genomes using machine learning and lineage-specificity

**DOI:** 10.1101/2021.03.19.436227

**Authors:** Munir Nur, Kelsey Wood, Richard Michelmore

## Abstract

Oomycete plant pathogens cause a wide variety of diseases, including late blight of potato, sudden oak death, and downy mildews of plants. These pathogens are major contributors to loss in numerous food crops. Oomycetes secrete effector proteins to manipulate their hosts to the advantage of the pathogen. Plants have evolved to recognize effectors, resulting in an evolutionary cycle of defense and counter-defense in plant–microbe interactions. This selective pressure results in highly diverse effector sequences that can be difficult to computationally identify using only sequence similarity. We developed a novel effector prediction tool, EffectorO, that uses two complementary approaches to predict effectors in oomycete pathogen genomes: (1) a machine learning-based pipeline that predicts effector probability based on the biochemical properties of the N-terminal amino acid sequence of a protein and (2) a pipeline based on lineage-specificity to find proteins that are unique to one species or genus, a sign of evolutionary divergence due to adaptation to the host. We tested EffectorO on *Bremia lactucae*, which causes lettuce downy mildew, and *Phytophthora infestans*, which causes late blight of potato and tomato, and predicted many novel effector candidates, while recovering the majority of known effector candidates. EffectorO will be useful for discovering novel families of oomycete effectors without relying on sequence similarity to known effectors.

## Introduction

Oomycetes are filamentous eukaryotic organisms that resemble fungi but are more closely related to brown algae (Baldauf 2003). Pathogenic oomycetes cause huge losses in agriculture and aquaculture worldwide (Derevnina et al. 2016; Haverkort et al. 2008). Oomycete pathogens and other parasites must continuously evade detection by their hosts in order to successfully infect and complete their life cycle. This is accomplished through the action of effectors, secreted proteins that interfere with host immune signaling and physiology to promote pathogen growth (Wang et al. 2019). To thwart the action of effectors and infection by the pathogen, plants have evolved receptors encoded by resistance genes (R genes) to detect effectors. Upon recognition, the receptors trigger a defense response that eventually results in localized cell death and disease resistance (Wang et al. 2019). Therefore, identification of effectors is important for both understanding how pathogens infect their hosts and for breeding disease resistance.

Most effector genes encode an N-terminal signal peptide that targets the protein for secretion outside of the pathogen and is cleaved from the mature peptide sequence. However, atypically secreted effectors lacking a signal peptide have also been observed in fungi and oomycetes (Liu et al. 2014), and atypical secretion has been documented in protozoans at the host–parasite interface (Balmer and Faso 2021). Through conventional or atypical secretion, all effectors are initially exported from the pathogen into the apoplast, where they either remain (in the case of apoplastic effectors) or are translocated into the plant cell (in the case of cytoplasmic effectors).

For some oomycetes, such as *Phytophthora* spp. and downy mildew pathogens, translocation of effectors has been linked to the N-terminal sequence directly following the signal peptide, which often contains one or more of several characteristic amino acid motifs. These include the RXLR and EER motifs (Birch et al. 2009; Rehmany et al. 2005; Whisson et al. 2007) and the LXLFLAK motif, associated with Crinkling and Necrosis (CRN) causing effectors (Amaro et al. 2017). The precise mechanism of translocation is the topic of some controversy (Ellis and Dodds 2011); regardless, these motifs have proven useful for effector prediction in multiple pathogen genomes. Many of these cytoplasmic effectors have been validated by testing the candidate proteins for recognition by resistant genotypes (avirulence activity) or immune suppression (virulence activity) *in planta* (Deb et al. 2018; Fabro et al. 2011; Pecrix et al. 2019; Pel et al. 2009; Stassen et al. 2013; Zheng et al. 2014).

While the RXLR motif is clearly a conserved effector motif in *Phytophthora* species, effectors in the related downy mildew pathogens show divergence and diversification of this motif (Bailey et al. 2011; Sharma et al. 2015). Due to the sequence divergence of the RXLR motif in downy mildew species, both string- and Hidden Markov Model (HMM)-based searches recover fewer effector candidates in these pathogens compared to *Phytophthora* species (Baxter et al. 2010; McGowan and Fitzpatrick 2017). Another approach to predict effectors in downy mildew species (as well as in *Phytophthora*) is to search for the structurally conserved WY domain, which is present in ∼25–50% of RXLR effectors in *Phytophthora* and downy mildew pathogens (Bos et al. 2010; Dou et al. 2008; Win et al. 2012) and has been implicated in effector function (He et al. 2019; King et al. 2014). While HMM searches for the WY domain have a low false positive rate for predicting candidate effectors (Wood et al. 2020), they likely have a high false negative rate because not all RXLR effectors have WY domains and, furthermore, there may be *bona fide* effectors with novel N-terminal motifs and uncharacterized effector domains.

Unlike oomycetes, fungi have no widely conserved effector motifs or domains that are present across all species. Degenerate RXLR-like motifs have been found in some fungal pathogens (Gu et al. 2011; Kale et al. 2010) and others have been found to have an N-terminal Y/F/WxC motif (Godfrey et al. 2010). EffectorP, a machine learning based approach to identify effectors in fungal genomes using biochemical characteristics, successfully expanded the pool of candidate effectors in fungal species (Sperschneider et al. 2016, 2018). Machine learning-based effector prediction pipelines, such as EffectorP, work by building effector classification models trained on known effectors and secreted non-effectors from the target organisms. Once a suitable model is determined, the classifier calculates the biochemical features from the amino acid sequence of the protein and outputs a probability that the protein is an effector based on those features. At the initial submission of this paper, machine learning had not been applied to effector prediction in oomycetes. More recently, a new version of EffectorP has been published (EffectorP 3.0), which has the ability to predict oomycete effectors as well as fungal effectors and additionally classifies them as either apoplastic or cytoplasmic (Sperschneider and Dodds 2022). EffectorP 3.0 uses an ensemble of machine learning classifiers trained on fungal and oomycete effectors to predict effectors starting from predicted secreted proteomes (Sperschneider and Dodds 2022). In this paper, we present a complementary machine learning effector prediction pipeline for oomycetes that gives similar accuracy as EffectorP 3.0 and may give higher specificity, especially in downy mildew species. We also present a pipeline to identify lineage-specific proteins that can be combined with the machine learning pipeline to filter for effector candidates.

Lineage specificity is often a characteristic of effectors due to the constant evolutionary cycle between pathogen and host. Pathogenicity-related proteins, such as effectors (as well as their corresponding R genes), often show increased rates of sequence divergence (Dodds et al. 2006; Dong et al. 2015). This is especially true for organisms that have narrow host ranges, such as downy mildew pathogens. For example, the genomes of the downy mildew species *Peronospora effusa* and *P. tabacina* are predicted to contain ∼1,800 and ∼3,000 species-specific genes, respectively (Fletcher et al. 2018). In addition, comparison between two isolates of *P. effusa* revealed >600 isolate-specific genes (Fletcher et al. 2018). These species- and isolate-specific genes may have arisen through co-adaptation to specific hosts or in response to different resistance genes of the same host, respectively. We reasoned that by using lineage-specificity in combination with a machine learning-based effector classifier, we would be able to expand the candidate effector repertoires of oomycete pathogens, especially those with narrow host ranges, as these approaches are able to find candidates that have diverged from effectors with known motifs and domains. For effector prediction, we define “lineage-specific” broadly to mean a sequence that is specific to one phylogenetic lineage: either at the genus, species, or isolate level. For the oomycetes studied in this paper, we classified proteins as lineage-specific at the species level if there were no close orthologs in any of the other sequenced oomycete genomes. The classification of a protein as lineage-specific depends greatly on what genomic data is available for other species, and thus this concept is a more of a heuristic tool for narrowing down lists of effector candidates by removing conserved proteins, rather than a biologically-relevant characteristic of any given protein.

Here we present EffectorO: a novel pipeline to predict effectors in oomycete genomes that avoids the limitations of motif/domain-based searches. EffectorO consists of a machine learning based effector classifier trained on a diverse set of oomycete effectors (EffectorO-ML) combined with a lineage-specific protein sequence identification pipeline (EffectorO-LSP). We used two economically important oomycete pathogens as model organisms to evaluate EffectorO: *Bremia lactucae*, an obligate biotrophic downy mildew pathogen with complete host-specificity to *Lactuca* spp., and *Phytophthora infestans*, which causes late blight of potato and tomato. EffectorO had high accuracy for classifying protein sequences as effectors, predicted thousands of novel effectors in these two oomycete genomes, and recovered known effectors. Effector prediction was then expanded to the genomes of an additional 26 oomycete species. One candidate effector from EffectorO was validated by demonstrating the elicitation of cell death on downy mildew resistant genotypes by an effector of *B. lactucae*. This candidate contained no obvious canonical sequence motifs (such as RXLR or EER). Thus, this paper shows that a machine learning approach, especially when combined with a lineage-specificity analysis, can be useful in gene annotation and effector prediction in oomycete genomes.

## Materials and Methods

### Positive and negative training sets

Our positive training set consisted of oomycete effectors shown experimentally to be either recognized by R genes (avirulence activity) and/or to have an immune-suppressing effect on the host (virulence activity) (Supplemental Table 1). Seventy-seven of these sequences were obtained from the Pathogen-Host Interaction database (PHI-base) (Urban et al. 2019) and 15 effector candidates were added from *B. lactucae* that had avirulence (Giesbers et al. 2017; Pelgrom et al. 2019; Stassen et al. 2013; Wood et al. 2020) or virulence activity (Wood et al. 2020), resulting in an initial set of 92 sequences. We then clustered sequences using CD-HIT at a sequence identity threshold of 70% and word length of five amino acids in order to identify and remove closely related sequences (Fu et al. 2012; Li and Godzik 2006). This resulted in a final positive training set of 88 unique and distinct oomycete effectors.

Our negative training set consisted of representative oomycete sequences that were predicted to be secreted and very highly conserved (thus, unlikely to be effectors) throughout the genomes of 28 oomycete species (Supplemental Table 2). To find these sequences, we first performed an Orthofinder search on the oomycete genomes at an e-value cutoff of 1e-10 and a query coverage cutoff of 90% (Emms and Kelly 2015, 2019). We then retrieved ortholog centroids, with the species chosen at random, that were present in at least 26 genomes. We predicted signal peptides using SignalP v4.1 (sensitive mode) (Petersen et al. 2011). To further reduce potential model bias and redundancy in the set of non-effectors, we clustered sequences using CD-HIT and removed sequences with greater than 40% identity. This identified 320 sequences, from which a training set of 88 sequences was chosen at random to obtain a balanced 1:1 positive to negative training set ratio.

### Feature selection and encoding for classical machine learning models

To calculate biochemical characteristics of effectors and secreted presumed non-effectors, we used published amino acid scales from ExPASyProtScale (ExPaSy; Expert Protein Analysis System 2004) and evaluated six features that could be informative for effector classification. We focused on the first 100 amino acids of the N-terminus of protein sequences, with the rationale that (1) secretion and translocation signals are typically encoded in the N-terminus and (2) effectors may differ in their C-terminal domains depending on their specific functions (for example, not all validated oomycete effectors have WY domains and RXLR effectors show high rates of positive selection in the C-terminus (Win et al. 2007)). To ensure the feature scales were equivalent to each other, we transformed the calculated feature distributions to zero mean and unit variance (Z-score). We calculated the average values for six N-terminal biochemical features: the grand average of hydropathy (GRAVY)(Kyte and Doolittle 1982), hydrophobicity (Fauchere and Pliska 1983), surface exposure (Janin 1979), disorder propensity (Dunker et al. 2001), bulkiness (Zimmerman et al. 1968), and residue interface propensity (Jones and Thornton 1997) of the first 100 amino acids using data from ExPASyProtScale (ExPaSy; Expert Protein Analysis System 2004).

PONDR VSL2 (Peng et al. 2006) was used as an additional method to visualize levels of intrinsic disorder for effectors and secreted non-effectors across the N-terminal sequence. The average sequence disorder was calculated at each amino acid position for the first 160 amino acids, including the signal peptide, in order to visualize the differences between candidate effectors and non-effectors.

### Machine learning architectures

For the classical machine learning pipeline, we tested various parametric and non-parametric models and evaluated what worked best for effector classification based on the N-terminal protein sequence. We tested and compared the following models using multiple cross-validation trials (resampling procedures that rotate training and testing splits): Random Forest, Naïve Bayes (Bernoulli and Gaussian), Logistic Regression, and K-Nearest Neighbors. To reduce overfitting, we monitored training and testing scores (accuracy, sensitivity, specificity, false positive rate, and area under the precision-recall curve). Models were used from the scikit-learn machine learning library in Python (Pedregosa et al. 2011). We compared Logistic Regression (default parameters), Naive-Bayes models (Gaussian and Bernoulli, default parameters), Random Forest (with 400 estimators and a max depth of 4), and the K-Nearest Neighbors classifier (with the neighbors parameter set to 5).

### Feature encoding for Convolutional Neural Network models

In order to account for signals that may be concealed through net average calculations, we also tried using a convolutional neural network (CNN) to extract positional-dependent differences in amino acid sequence between effectors and non-effectors. We used the first 100 amino acid values for each feature and padded the ends with zeros for sequences shorter than 100 aa. For each protein in our positive and negative training sets, this feature encoding resulted in a 6 x 100 matrix to include every positional encoding for all six biochemical features in the first 100 amino acids.

### Genome data and secretome prediction

Genomic sequences from 28 oomycete species were used to test EffectorO (list of species and citations in Supplemental Table 2). We then predicted open reading frames (ORFs) for all genomes. ORFs were used instead of gene annotations for several reasons: (1) effector genes often consist of a single exon; (2) effector genes are often missed by gene annotation programs; and (3) to account for differences in gene annotation approaches used in different genomes. We predicted ORFs from each genome using BioPerl’s “translate_strict” module, using the canonical start codon (ATG), a maximum percent X of 0, and a minimum nucleotide length of 240 (Stajich et al. 2002). From translated ORFs, we then used SignalP v4.1 (sensitive mode) (Petersen et al. 2011) to predict secreted proteins. These predicted secreted proteins were then used in subsequent EffectorO-ML analysis and for the lineage specificity pipeline.

### Lineage-specificity

Proteins encoded by lineage-specific genes were obtained using the predicted secretome of a species as a BLASTp query against the translated open reading frames (ORFs) of other oomycete species (Supplemental Table 2). Conserved proteins were defined as proteins with a BLASTp hit in another species with an e-value cutoff of <10e-7, percent identity of >30%, and a query coverage of >30%. Similar BLASTp cutoffs are used in other pipelines to obtain species-specific genes for numerous species, including other oomycetes (Rujirawat et al. 2019; Zhou et al. 2015). Protein sequences that were below these thresholds in another oomycete species were defined as lineage-specific. ORFs were used instead of gene models for the reasons described above.

#### Prediction of amino acid motifs and protein domains

To predict the RXLR-EER motif, we used a string search for [RQGH]XLR, [DE][DE][KR] for EER, and HMM for RXLR-EER developed by Whisson et al. (2007) for *Phytophthora* species. To predict WY domains, we used an HMM built from a training set of WY domain proteins from three *Phytophthora* species by Boutemy et al. (2011). To predict CRNs, we used an HMM built from sequences of CRNs from *Phytophthora* species (Fletcher et al. 2018). HMMer v3.1 was used for all HMM searches (Eddy 2011). Default settings were used for HMMer, except we used a bitscore cutoff of >0 to classify WY domain proteins (as suggested by Boutemy et al. (2011)) and CRNs, which is more inclusive than the default cutoff. To predict enzymatic and other characterized functional domains, the amino acid sequences of EffectorO candidates were queried for Pfam predicted protein domains using the HMMER website (https://www.ebi.ac.uk/Tools/hmmer/search/hmmscan). ProfileHMM search was performed with an e-value cutoff of 0.1 against the Pfam database (Mistry et al. 2020). Resulting Pfam predictions were checked for accuracy of annotation and were manually categorized into domain families.

#### Transcriptome data

Transcriptome data for *B. lactucae* were downloaded from the NCBI short read archive (NCBI BioProject PRJNA523226). These reads represent Illumina HiSeq 4000 sequencing from RNA obtained from *B. lactucae* infecting 7-day-old lettuce seedlings and represent the full lifecycle of *B. lactucae* from inoculation to sporulation (timepoints: 0h, 3h, 6h, 12h 24h, 3d, 5d, 7d). Read counts for all predicted ORFs were generated using samtools (Li et al. 2009); resulting read counts per ORF were then normalized by ORF length and log-normalized for visualization.

#### Sequence analysis and protein structural prediction

To search for homologous proteins and compare sequences, protein BLAST (BLASTp) was used against the NCBI database (https://blast.ncbi.nlm.nih.gov/Blast.cgi) (Sayers et al. 2020) with the amino acid sequence of the protein of interest as the query. ClustalW was used to align amino acid sequences for comparison (Thompson et al. 1994). Protein structures were predicted using the full amino acid sequences (including the predicted signal peptide) using RoseTTAfold on the Robetta web server with default settings (https://robetta.bakerlab.org/)(Baek et al. 2021).

#### Scripts

All scripts for running EffectorO-ML and EffectorO-LSP can be found on Github at https://github.com/mjnur/oomycete-effector-prediction. A web app for EffectorO-ML is available at https://bremia.ucdavis.edu/effectorO.php.

### Candidate gene cloning and agroinfiltration

The effector candidate *BLE01* was cloned from *B. lactucae* genomic DNA using Gateway cloning into pEG100 (35S promoter) as described previously (Wood et al. 2020). The signal peptide was excluded from the coding sequence to ensure expression inside the plant cell. Two distinct alleles were obtained. *Agrobacterium tumefaciens* strain C58Rif+ was transformed with pEG100-BLE01 and infiltrations were performed on 3- to 5-week-old lettuce plants as described previously (Wood et al. 2020). Leaves were scored for necrosis 4–5 days after infiltration. *A. tumefaciens* containing pEG100 (empty vector) and pEG100-GFP were used as negative controls; *A. tumefaciens* containing pBAV139-35S:HopM1 and pBAV139-35S:AvrPto were used as positive controls for induction of cell death (Wroblewski et al. 2009).

## Results

### Effector and non-effector proteins differ in their biochemical characteristics

To determine whether various N-terminal amino acid characteristics would be sufficient to distinguish effectors from non-effector proteins, we compared predicted biochemical parameters between validated oomycete effectors and secreted, conserved proteins (presumed non-effectors). We found that validated effectors had higher intrinsic disorder and had lower scores for GRAVY, hydrophobicity, exposed residues, bulkiness, and interface, compared with secreted, conserved proteins (Fig. 1A). The intrinsic disorder score was significantly higher in validated effectors, starting from just after the signal peptide (first ∼20 aa) until approximately 80 aa after the start codon (Fig. 1B).

**Figure 1.**
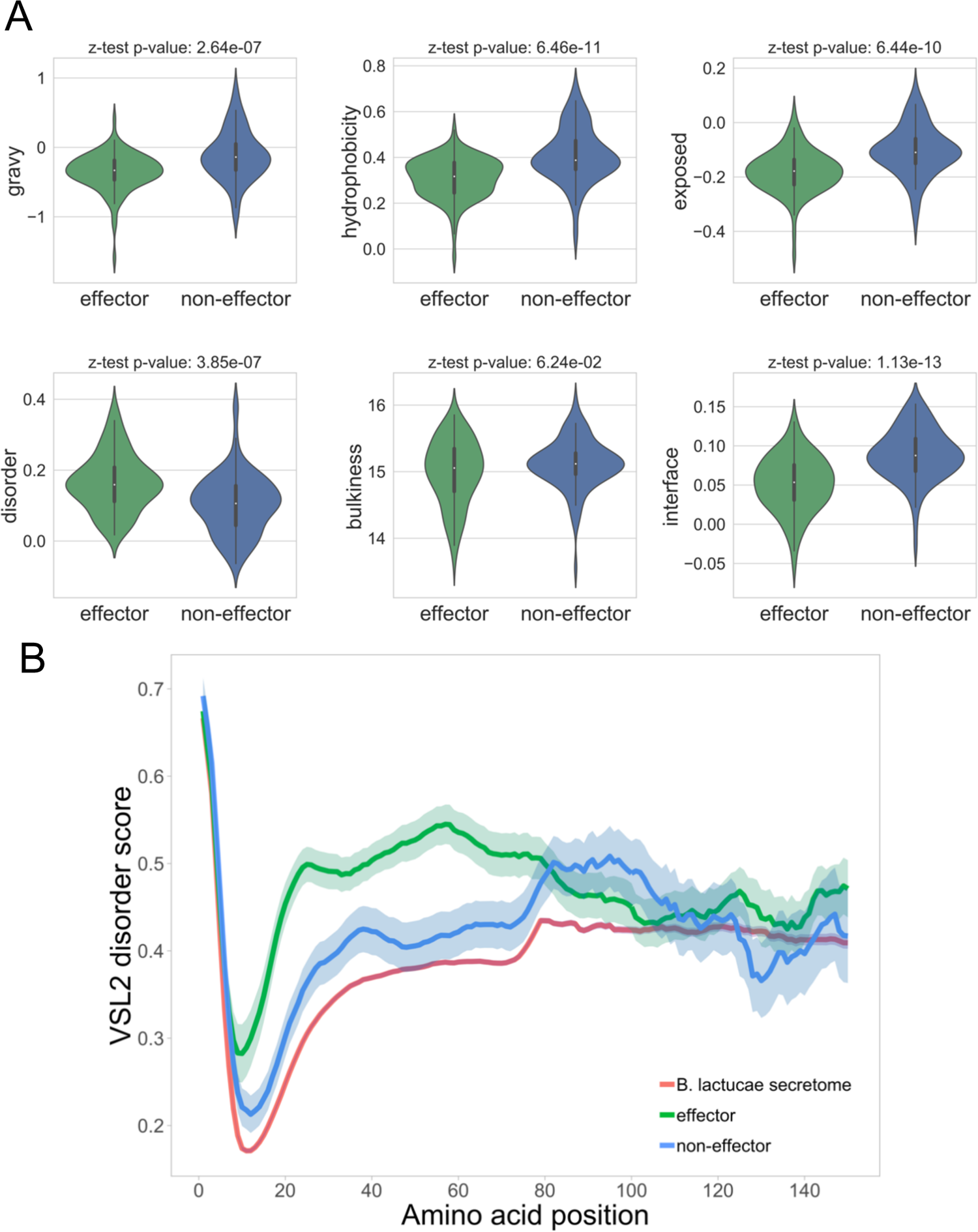
**(A)** Amino acid biochemical characteristics for validated oomycete effectors and secreted non-effectors as calculated by the ExPASyProtScale and used as features for the machine learning classifier. Violin plot shows the distribution of the N-terminal Z scores for each group of proteins, with the p-value shown above the plot. **(B)** Intrinsic disorder as calculated by PONDR VSL2 for validated oomycete effectors and secreted non-effectors with the *B. lactucae* secretome for reference. The average and standard error are shown for each group of proteins at each amino acid position.

### Evaluation of machine learning effector classifier models

To find the effector classifier with the best performance, we evaluated six machine learning models (Random Forest, Linear SVCs, Bernoulli and Gaussian Naïve Bayes, Logistic Regression, and K-Nearest Neighbors) as well as a convolutional neural network model that takes into account positional information for classification. The Random Forest classifier initially outperformed other models, so we attempted several combinations of parameters, adjusting the number of estimators (n_estimators) and maximum tree depth (max_depth), while monitoring the cross-validation accuracy. The best receiver-operator curve (ROC) was achieved by the Random Forest classifier (with parameters n_estimators = 400 and max_depth = 4) with an area under the curve (AUC) of 0.89 (Figure 2) and an overall 5-fold cross-validation accuracy of 84.26% (Table 1). The convolutional neural network was found to perform similarly but not better than the most accurate Random Forest model. Therefore, we used the Random Forest model for our EffectorO-ML pipeline to classify effectors based on the biochemical characteristics of the first 100 amino acids.

**Figure 2.**
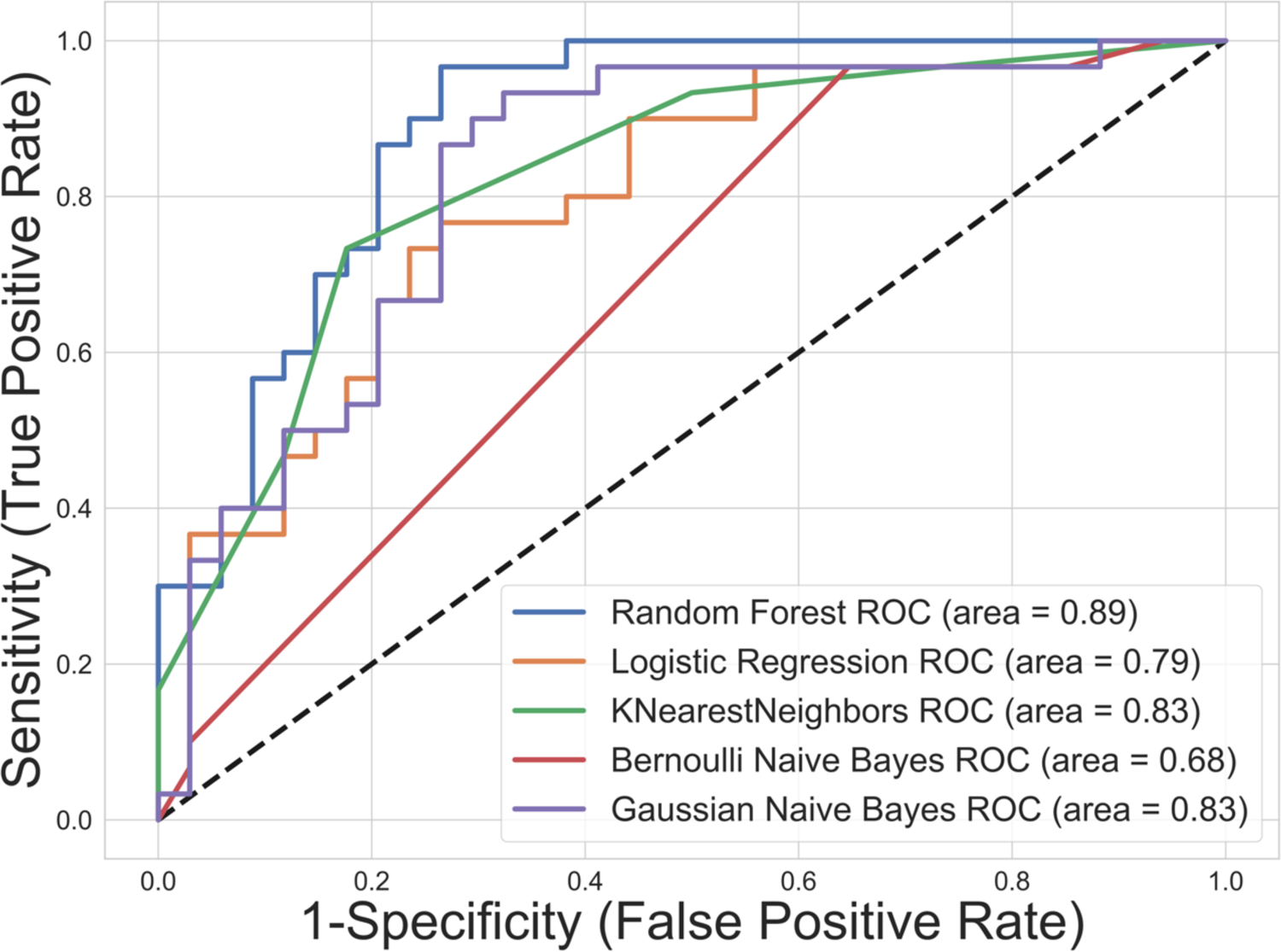
Receiver-operator curves (ROCs) for the various machine learning models tested on a cross validation fold using 1:1 effector to non-effector testing and training sets.

**Table 1.**
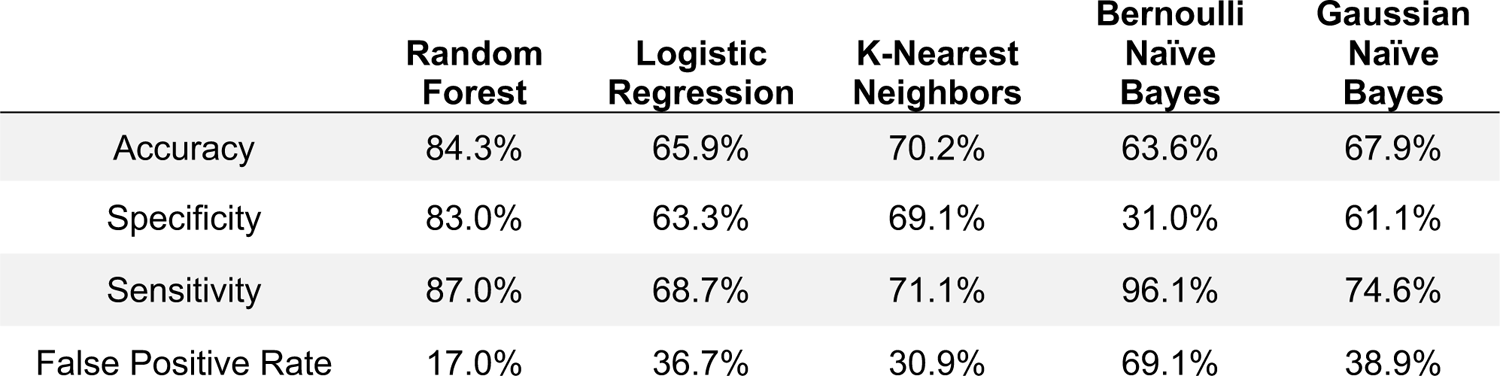
Average cross validation metrics for various machine learning models for classifying proteins as effectors or non-effectors

The Random Forest effector classifier uses a default probability threshold of 0.5 to classify a protein as an effector; this probability threshold can be modified depending on preference for either inclusivity/sensitivity or specificity. For the top performing Random Forest model, we calculated and averaged accuracy, specificity, sensitivity (true positive rate), and false positive rate along the five cross-validation folds for different probability thresholds, illustrating the tradeoffs between rates of false positives and true positives when varying probability thresholds (Table 2). To obtain another estimate of the false positive and false negative rates, we ran EffectorO-ML on a set of 634 predicted secreted proteins that were highly conserved (presumed non-effectors) in at least 24 of the 28 sequenced oomycete species used for EffectorO-LSP, as well as on the 41 secreted *B. lactucae* WY effectors (presumed true effectors). We found that a probability cutoff of 0.5 resulted in approximately 28% of these conserved proteins being classified as effectors while capturing 90% of the WY effectors (Figure 3 and Table 3). At a probability cutoff of 0.6, only 16% of the conserved proteins were classified as effectors, with 85% of the WY proteins classified as effectors (Table 3). Since we presume that most of the conserved secreted proteins are not effectors due to lack of lineage-specificity, 16% would be a good estimate of the false positive rate at the 0.6 threshold, which is similar to the false positive rate for string searches for the RXLR motif (Wood et al. 2020). The false positive rate can be decreased further if desired by increasing the probability threshold, but at the loss of capturing true effectors.

**Figure 3.**
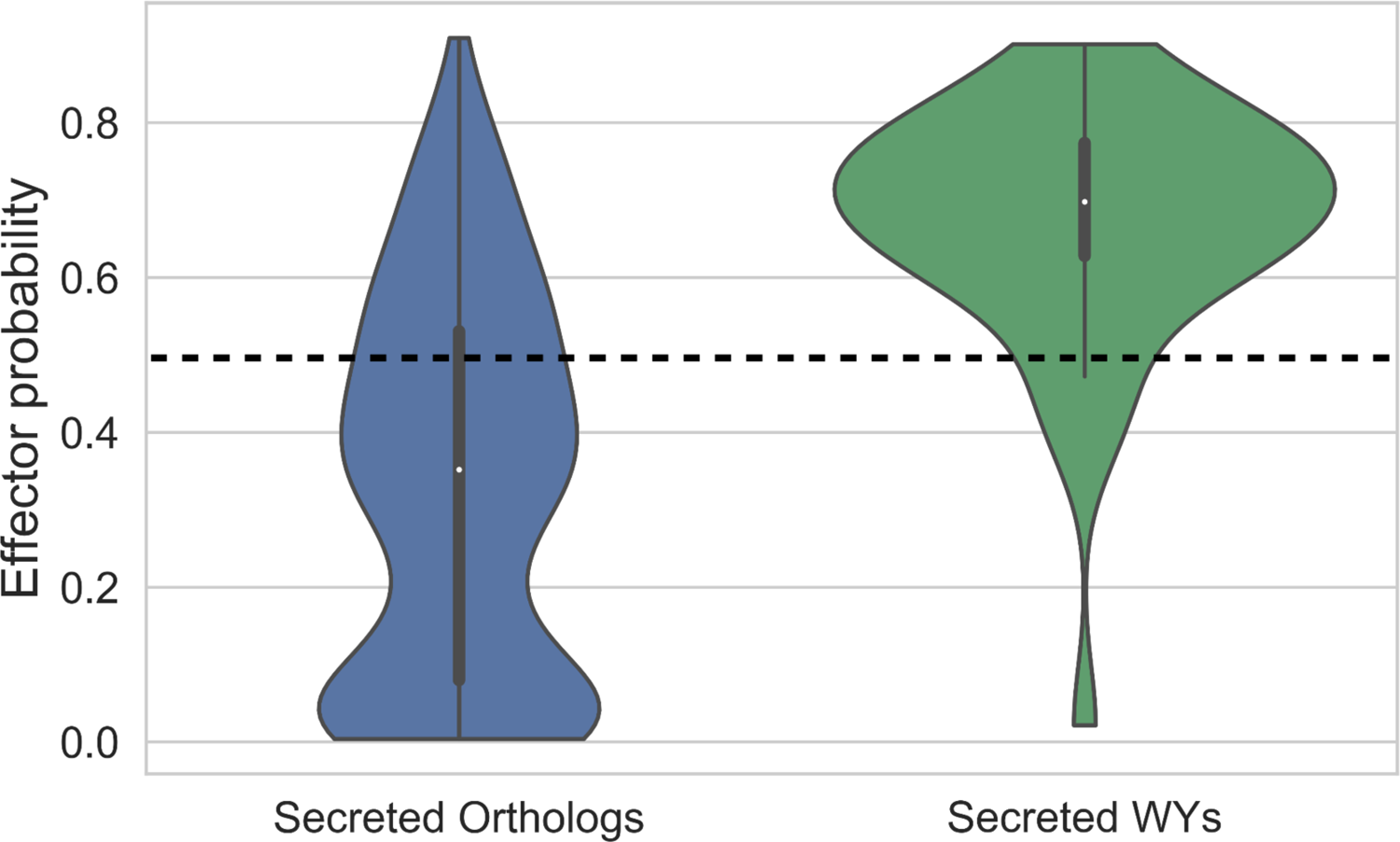
Violin plots of effector probabilities for conserved secreted oomycete orthologs and secreted *B. lactucae* WY domain containing proteins, generated by the top-performing Random Forest model. Dotted line at 0.5 shows the suggested inclusive cutoff for effector classification.

**Table 2.** Metrics at different probability thresholds for the most accurate Random Forest model

|  | Random Forest Model Effector Probability Threshold = |  |  |
| --- | --- | --- | --- |
|  | 0.5 | 0.6 | 0.7 |
| Accuracy | 84.3% | 79.0% | 76.2% |
| Specificity | 83.0% | 85.1% | 89.8% |
| Sensitivity | 87.0% | 73.0% | 62.7% |
| False Positive Rate | 17.0% | 14.9% | 10.2% |

**Table 3.** Number and percent of oomycete conserved secreted orthologous proteins (634 total) and *B. lactucae* secreted WY proteins (41 total) classified as an effector by the EffectorO-ML Random Forest model at different effector probability thresholds

|  | Random Forest Model Effector Probability Threshold = |  |  |  |  |
| --- | --- | --- | --- | --- | --- |
|  | 0.5 | 0.6 | 0.7 | 0.7 | 0.9 |
| Secreted Orthologs | 180 (28%) | 103 (16%) | 54 (8.5%) | 12 (1.9%) | 2 (0.3%) |
| Secreted WY Proteins | 36 (90%) | 34 (85%) | 20 (50%) | 5 (12.5%) | 1 (2.5%) |

### EffectorO predicts novel effectors in oomycete species and recovers known effectors

To find novel effectors from *P. infestans* and *B. lactucae*, we used our EffectorO-ML and LSP pipelines on the predicted secreted ORFs of these genomes. For *P. infestans*, we found 8,685 proteins (41% of the secretome) to be lineage-specific and 5,814 proteins (30%) to be classified as candidate effectors by machine learning, with 1,235 proteins (6%) present in both categories (Figure 4). For *B. lactucae,* we found 2,478 proteins (34% of the secretome) to be lineage-specific and 1,777 proteins (24%) to be classified as candidate effectors by machine learning, with 597 proteins (8%) present in both categories (Figure 4). EffectorO was largely able to recover proteins containing known effector signatures, such as the RXLR and EER motifs and the WY domain (Figure 4), which were not used as classifiers in our analysis but were contained in some proteins in the positive training set.

**Figure 4.**
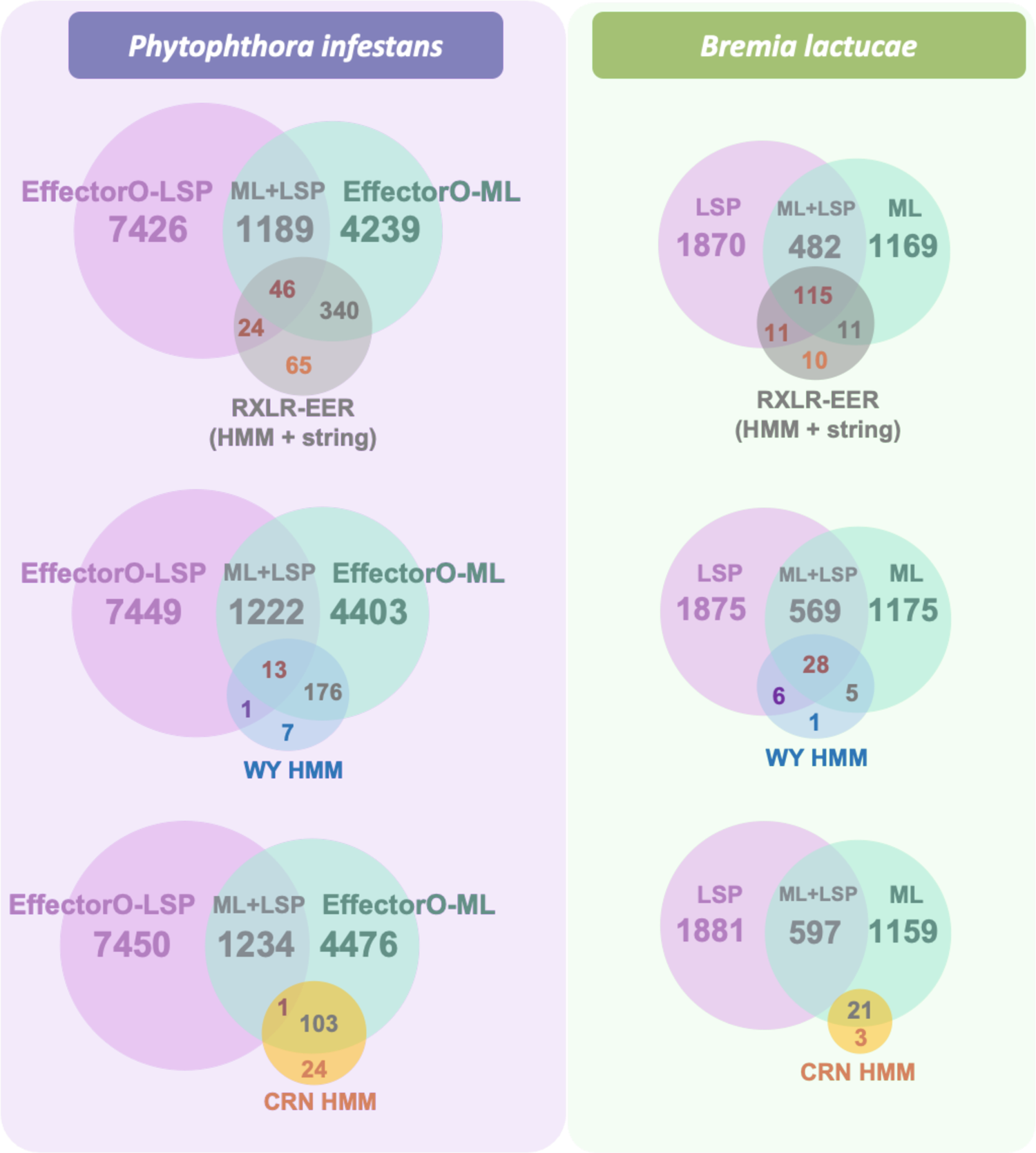
Venn diagrams showing the overlap between EffectorO predictions and canonical RXLR, WY, and CRN effector predictions for *P. infestans* and *B. lactucae*. Numbers from the predicted secretomes that are lineage-specific proteins (EffectorO-LSP) and/or predicted to be an effector by the machine learning classifier (EffectorO-ML, at probability score threshold of 0.5). RXLR-EERs were predicted using HMM and string search. WY domain proteins and CRNs were predicted using HMMs. Circle sizes are relatively proportional for each set but not precisely to scale.

### The majority of EffectorO candidates are expressed in B. lactucae

To investigate whether EffectorO candidates are expressed at the transcript level, we used an available RNA-seq dataset of *B. lactucae* grown on lettuce seedlings from 0 to 7 dpi (from inoculation to sporulation). We found that 73% of transcripts that encoded proteins predicted to be effectors by EffectorO-ML and EffectorO-LSP were expressed. The EffectorO-ML+LSP candidates had similar mean expression levels as transcripts encoding proteins containing WY domains and slightly lower expression than those containing RXLR-EER motifs (Figure 5). Transcripts that encoded proteins predicted to be effectors by either the ML or LSP pipeline, but not both, had lower mean expression levels and more proteins with zero reads present in the RNA-seq dataset (Figure 5). The full secretome had the lowest mean expression and also had 1,716 predicted transcripts with zero reads, suggesting that many of the predicted ORFs may not be expressed (Figure 5).

**Figure 5.**
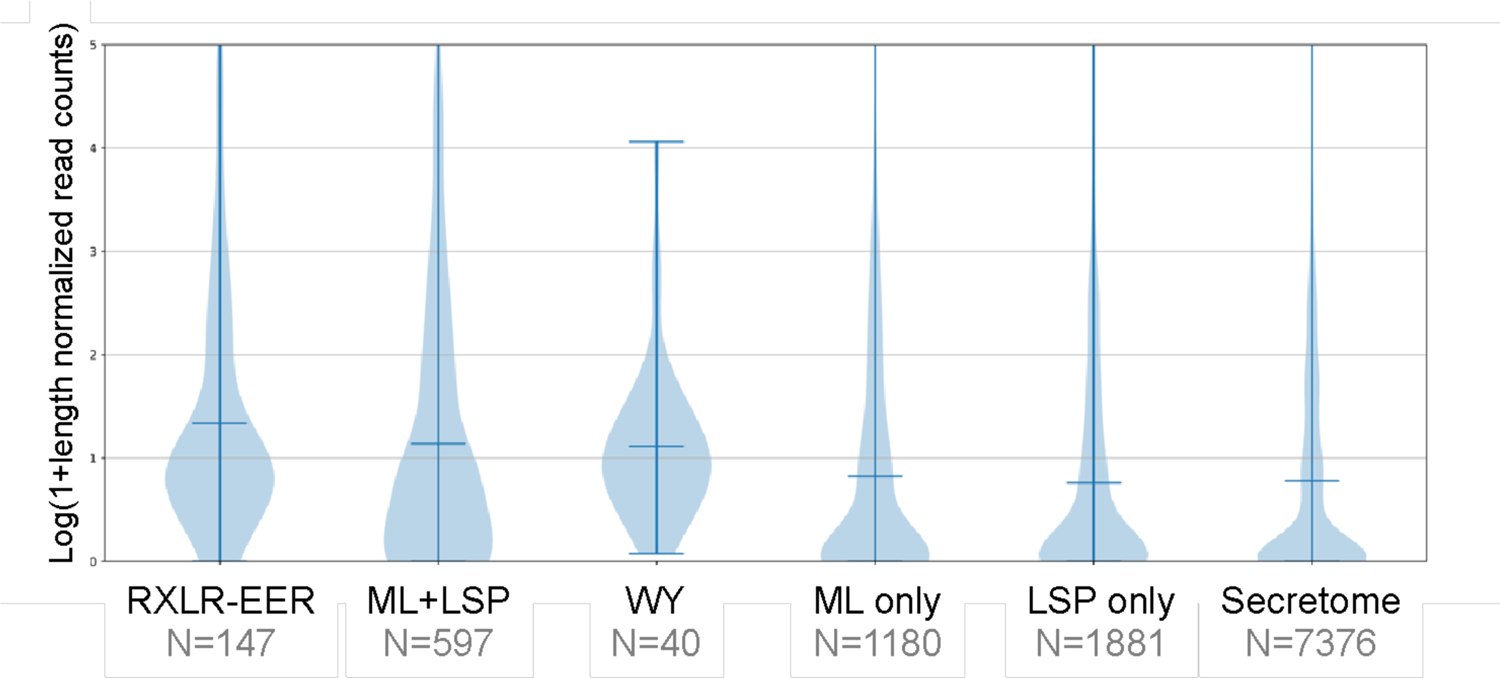
RNA-seq expression levels in *B. lactucae* (log of length-normalized read counts) for candidate effector transcripts encoding proteins with the RXLR-EER motif (RXLR-EER), WY domain (WY), EffectorO-ML-LSP (ML and LSP), EffectorO-ML only (ML only), EffectorO-LSP only (LSP only), or the full secretome. RNA was derived from a time course of *B. lactucae* infecting lettuce from 0–7 dpi. The number of ORFs in each category is given below the name.

#### The majority of EffectorO-ML candidates have no predicted functional domains

In order to investigate whether known functional domains were predicted to be present in the EffectorO-ML effector candidates, we queried EffectorO-ML candidates (with a score of 0.5 or higher) against the Pfam protein domain database. Most EffectorO candidates (88%) had no predicted Pfam domains (Table 4). There were 205 proteins that contained a total of 246 Pfam domains (with some proteins predicted to have multiple domains). The largest class of functional domains predicted was reverse transcription, or transposon-related domains, suggesting that these candidates may actually be pseudogenes disrupted by an insertion event (Table 5). Of the remaining EffectorO predictions with non-transposon-related domains, the most common domains predicted were enzymatic (including cell-wall degrading), protease, protein kinase, histidine phosphatase, lectin, cysteine-rich chitin binding, glycosyl hydrolase, and necrosis inducing protein domains (Table 5). All predictions are given in Supplemental Table 3.

**Table 4.** Pfam domains in *B. lactucae* EffectorO-ML candidates

| Category | Number of proteins |
| --- | --- |
| 1572 | No Pfam domains |
| 205 | With Pfam domains |
| **1777** | **Total EffectorO-ML candidates** |

**Table 5.** Top Pfam domain classifications for EffectorO-ML candidates (with at least six domains represented per family)

| Domain classification | # of Domains |
| --- | --- |
| **RT related** | **61** |
| Reverse Transcriptase | 39 |
| RNase H-like domain found in reverse transcriptase | 13 |
| LTR copia-type | 7 |
| Integrase zinc binding domain | 2 |
| **Enzymatic** | **35** |
| Carbohydrate metabolism and/or Cell-wall degrading | 24 |
| Other activity | 11 |
| **Protease** | **17** |
| **Protein kinase domain** | **15** |
| **Histidine phosphatase** | **13** |
| **Lectin** | **10** |
| **Cysteine-rich chitin binding** | **9** |
| **Chaperone** | **9** |
| **Necrosis inducing protein** | **8** |
| **Kazal-type serine protease inhibitor** | **7** |
| **Calmodulin-binding motif** | **7** |
| **Ribosomal protein** | **6** |
| **Elicitin** | **6** |

### EffectorO predicts candidate effectors in diverse oomycete pathogens

To determine whether EffectorO could predict effectors in other pathogens across the oomycete phylum, we tested EffectorO-ML and -LSP on a total of 28 published oomycete genomes. We found that even though EffectorO was trained primarily on effectors from downy mildew and *Phytophthora* pathogens, the algorithm predicted hundreds to thousands of candidate effectors in distantly related oomycetes infecting animals (e.g., *Achlya hypogyna*, *Aphanomyces* spp., *Saprolegnia* spp.), as well as oomycetes from the genera *Albugo* and *Pythium* (Figure 6). Critically, EffectorO-ML predicted a similar number of effectors in the free-living oomycete species *Thraustotheca clavata* (14.6% of the genome) as the false positive rate for EffectorO (17.0%, Table 1).

**Figure 6.**
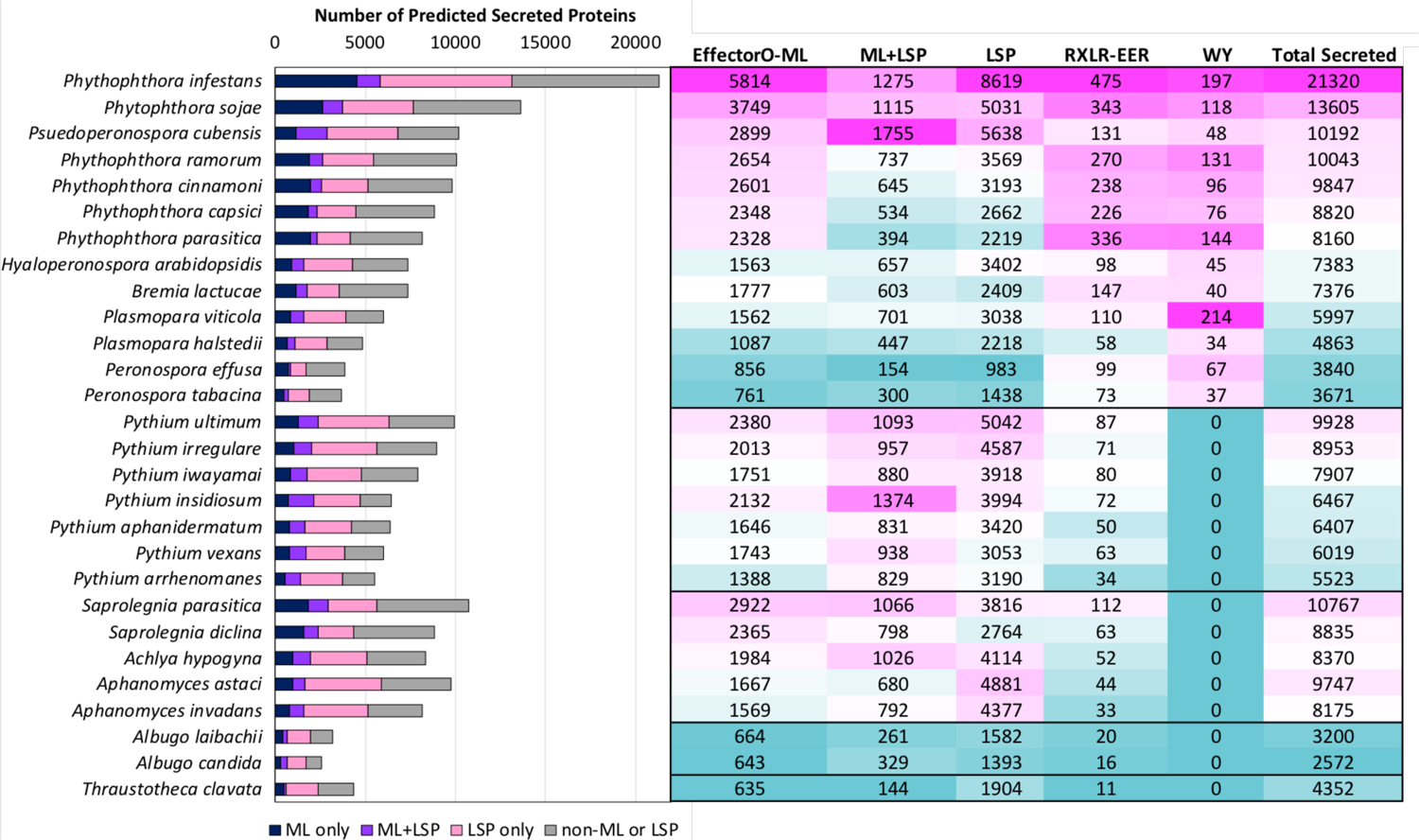
Number of predicted secreted proteins (translated ORFs) classified as effectors using EffectorO-ML only (cutoff = 0.5) (dark blue bars), EffectorO-LSP only (pink bars), EffectorO-ML and LSP (purple bars), and those classified as non-effectors (grey bars) in 28 species of oomycetes, including Peronosporales (*Phytophthora* and downy mildew species), *Pythium* species, animal pathogens (*Achlya, Saprolegnia*, and *Aphanomyces* spp.), *Albugo* spp., and a free-living oomycete (*Thraustotheca clavata*).

We compared effector predictions from EffectorO-ML for the 28 oomycete genomes to EffectorP 3.0 predictions. We found that EffectorP 3.0 predicts larger numbers of effectors in each genome (Table 6), with about 42% overlapping between the sets of predictions. For example, for *B. lactucae*, EffectorP 3.0 predicts 3111/7376 (42.2%) proteins in the secretome as effectors, while EffectorO predicts 1777/7376 (24.0%). In the free-living oomycete *T. clavata*, EffectorP 3.0 predicts 1918/4352 (44.1%) as effectors, while EffectorO predicts 635/4352 (14.6%).

**Table 6.** Number of effectors in 28 oomycete genomes predicted by EffectorO-ML and EffectorP 3.0.

| Species | Unique to EffectorO-ML | Predicted by both | Unique to EffectorP 3.0 |
| --- | --- | --- | --- |
| *Achlya hypogyna* | 1210 | 774 | 1317 |
| *Albugo candida* | 195 | 448 | 753 |
| *Albugo laibachii* | 177 | 487 | 953 |
| *Aphanomyces astaci* | 886 | 781 | 1945 |
| *Aphanomyces invadans* | 797 | 772 | 1456 |
| *Bremia lactucae* | 618 | 1159 | 1945 |
| *Hyaloperonospora arabidopsidis* | 582 | 981 | 1694 |
| *Peronospora effusa* | 315 | 541 | 858 |
| *Peronospora tabacina* | 254 | 507 | 953 |
| *Phytophthora capsici* | 1058 | 1290 | 1597 |
| *Phytophthora cinnamoni* | 1313 | 1288 | 1739 |
| *Phytophthora infestans* | 2841 | 2973 | 4080 |
| *Phytophthora parasitica* | 945 | 1383 | 1548 |
| *Phytophthora ramorum* | 1303 | 1351 | 1744 |
| *Phytophthora sojae* | 1936 | 1813 | 2183 |
| *Plasmopara halstedii* | 340 | 747 | 1297 |
| *Plasmopara viticola* | 498 | 1064 | 1521 |
| *Psuedoperonospora cubensis* | 1316 | 1583 | 1618 |
| *Pythium aphanidermatum* | 884 | 762 | 1039 |
| *Pythium arrhenomanes* | 791 | 597 | 814 |
| *Pythium insidiosum* | 1152 | 980 | 1069 |
| *Pythium irregulare* | 884 | 1129 | 1926 |
| *Pythium iwayamai* | 843 | 908 | 1477 |
| *Pythium ultimum* | 1111 | 1269 | 2043 |
| *Pythium vexans* | 1039 | 704 | 901 |
| *Saprolegnia diclina* | 1367 | 998 | 1236 |
| *Saprolegnia parasitica* | 1664 | 1258 | 1484 |
| *Thraustotheca clavata* | 163 | 472 | 1444 |

### An EffectorO predicted effector candidate causes cell death when expressed in lettuce

To test whether any EffectorO candidates were recognized effectors (i.e., have avirulence activity), we cloned a gene from *B. lactucae* that was within the mapping region for *Avr6,* as determined by sequence-based mapping of segregating sexual progeny of *B. lactucae* isolates (R.J. Gil, unpublished data). This region contained no genes encoding RXLR or EER motifs nor WY domains, but did contain one ORF encoding a protein that was classified as a probable effector by both EffectorO-ML and Effector-LSP. The ORF encoding this protein was not found in the gene models (i.e., was not predicted to be a gene by the annotation software). The candidate effector, named BLE01 for (*<u>B</u>remia lactucae* <u>E</u>ffectorO candidate <u>1</u>), caused cell death when transiently expressed in the lettuce cultivar Sabine, which contains *Dm6*, but not Cobham Green, which does not contain any known *Dm* genes (Fig 7A). When we screened a larger panel of lettuce cultivars with various *Dm* genes, we found that it also elicited cell death in RYZ2164, but not in the other seven cultivars with other *Dm* genes or in the non-host plant *Nicotiana benthamiana* (Fig 7B).

**Figure 7.**
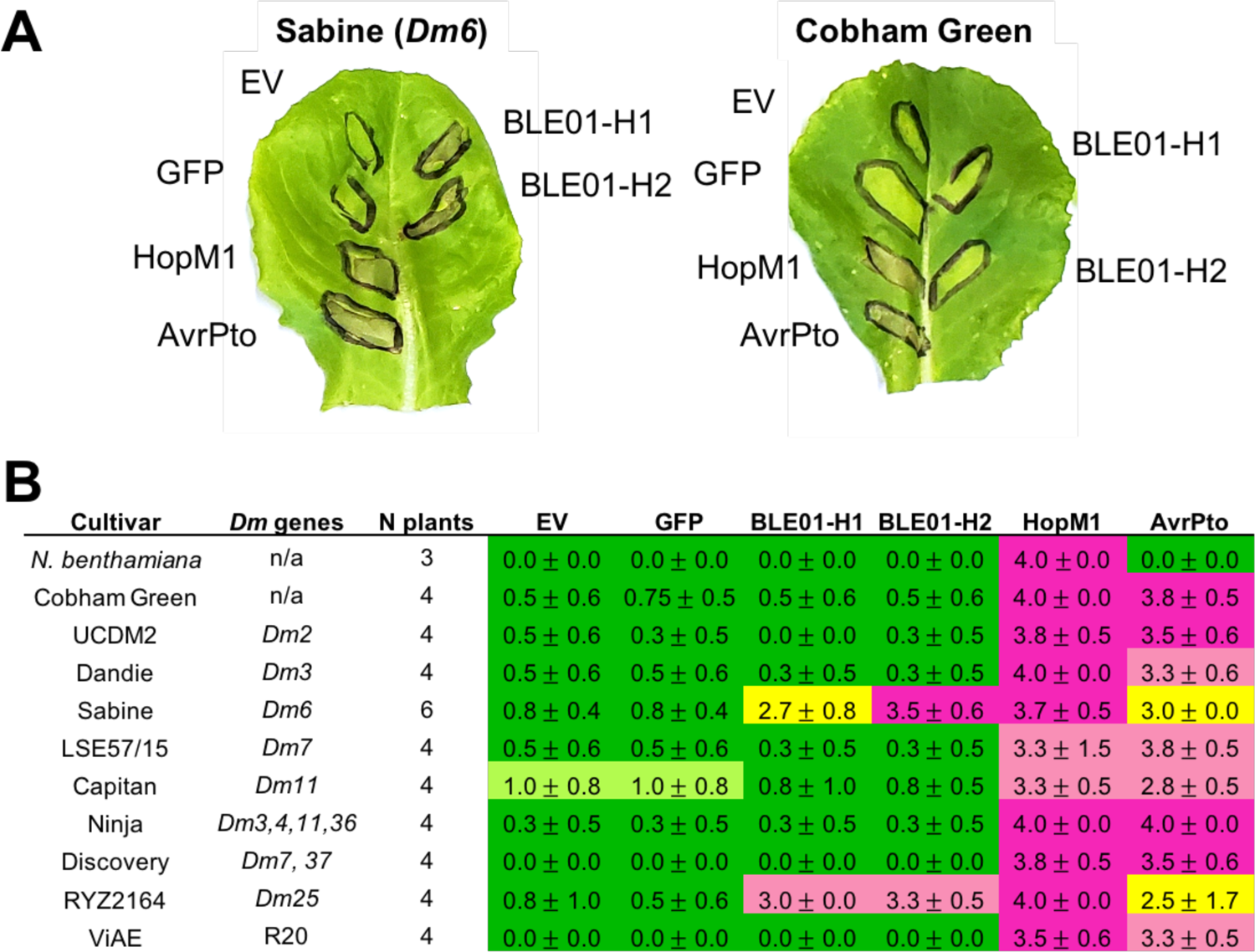
(A) Example transient expression results from lettuce infiltrated with *A. tumefaciens* containing two candidate effector alleles from *B. lactucae* (BLE01-H1 and BLE01-H2); the negative controls, empty vector (EV) and green fluorescent protein (GFP); and the positive controls for cell death, *Pseudomonas syringae* pv. *tomato* effectors HopM1 and AvrPto, which cause strong and medium necrosis in lettuce, respectively. Leaves are representative of four replicates and photos were taken 4 days after infiltration. (B) Necrosis scores four days post-agroinfiltration of the candidate effectors and controls in a panel of lettuce cultivars and the non-host *Nicotiana benthamiana*. Necrosis was scored as follows: 0 = no necrosis or chlorosis, 1 = mild chlorosis, 2 = chlorosis, 3 = chlorosis and necrosis, 4 = full necrosis. Number of replicates (N), averages, and standard deviations are shown for each line and effector combination.

#### BLE01 is similar in sequence and structure to WY domain proteins

To investigate potential protein domains of BLE01, the amino acid sequence of the protein was used as a BLAST query against the *B. lactucae* genome. One of the top BLAST hits was a previously characterized WY domain containing protein, BSW13 (Wood et al. 2020). There was 61% amino acid identity in the overlapping region, which consists of the entire amino acid sequence for BSW13 and the N-terminal half of the sequence for BLE01 (Figure 8). Since BSW13 was predicted to contain a WY domain, we performed a search using the HMM for the WY domain on the BLE01 sequence to see if there might be a low scoring WY domain predicted for BLE01 that had been previously missed. We found that BLE01 was predicted to have a low scoring WY domain in the same region as BSW13, with a domain e-value of 0.029 and a bitscore of 1.8. We discovered that this protein had been originally filtered out from our HMM search for WYs by the default bias filter in HMMer.

**Figure 8.**
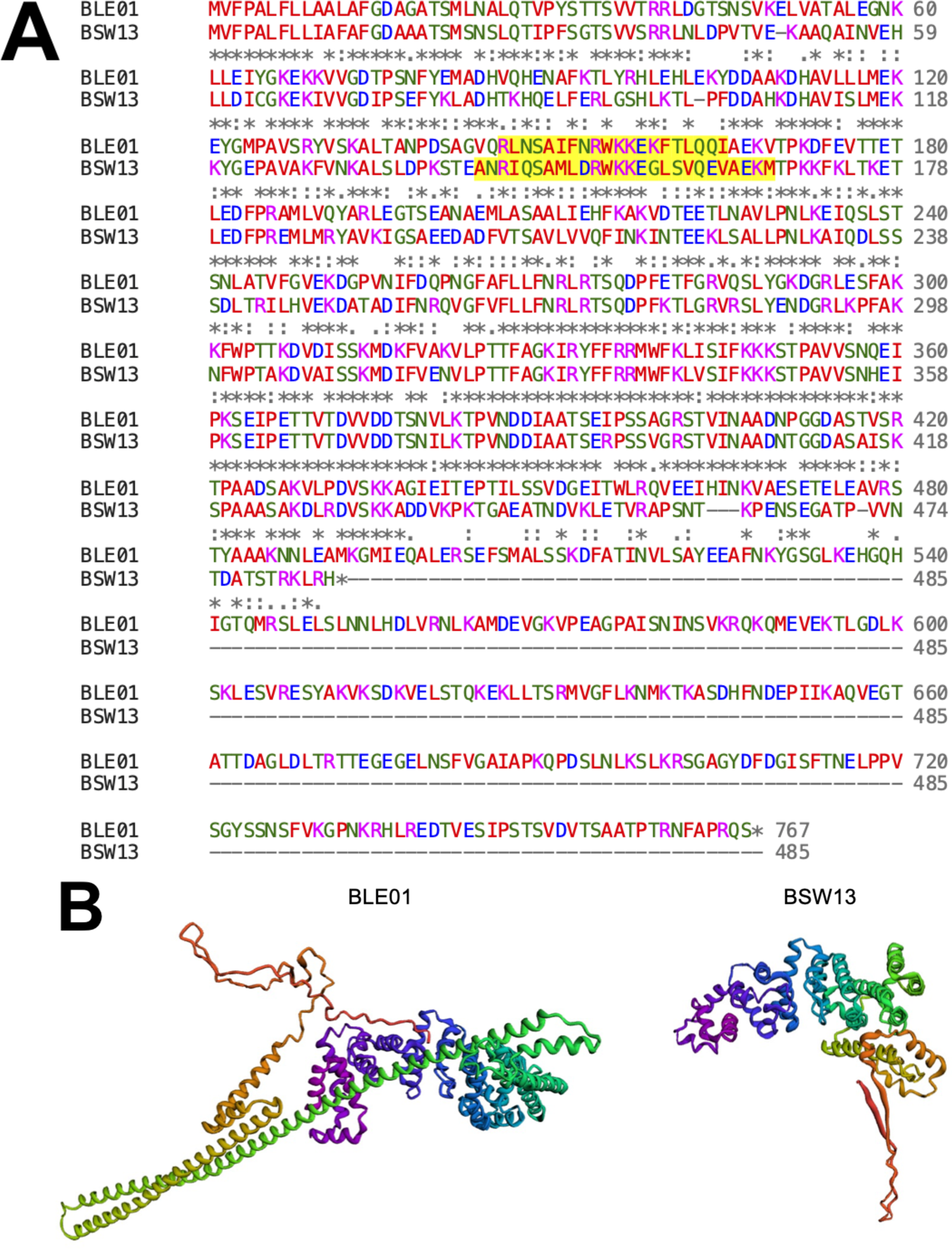
(A) Pairwise alignment between amino acid sequences of BLE01 and BSW13. Amino acids are colored red for hydrophobic, blue for negatively charged, magenta for positively charged, and green for all other amino acids. Similarity is indicated below the alignment with an asterisk (*) for matching, a colon (:) for strongly similar, and a period (.) for weakly similar amino acids. The predicted WY domains for each protein are highlighted in yellow. (B) Protein structure predictions from RoseTTAfold for BLE01 and BSW13. The protein structure is rainbow color coded from N-terminus (purple) to C-terminus (red).

To predict three-dimensional protein structures for BLE01 and BSW13, we used the RoseTTAfold web server (https://robetta.bakerlab.org/). Both proteins were predicted to have six alpha helical bundles (Figure 8B), which resemble the repeated WY domains as seen in other effector proteins that have been characterized by x-ray crystallography (Chou et al. 2011; Boutemy et al. 2011; Leonelli et al. 2011; Win et al. 2012; He et al. 2019).

## Discussion

We have demonstrated that our EffectorO pipeline can predict novel candidate effectors using a machine learning-based approach combined with a lineage-specificity criterion for oomycete effector classification. EffectorO can be used to mine oomycete genomes to expand the set of predicted effectors without relying on canonical N-terminal motifs (such as RXLR, EER, and LXLFLAK) or C-terminal domains (such as the WY domain); thus, it can reveal effectors that were previously missed by only searching for these features. EffectorO predicts an additional ∼600 ML+LSP candidate effectors from *B. lactucae* and ∼1,200 additional ML+LSP candidate effectors from *P. infestans*. The genome of *P. infestans is* larger than the genome of *B. lactucae (150 Mb vs 240 Mb)*, which may in part explain why more effectors were predicted in that genome (as well as in other *Phytophthora* species). The percentage of effectors predicted in the secretome for both species by EffectorO-ML was approximately 25%, which is similar to what was predicted for fungal secretomes with EffectorP (Sperschneider et al. 2018). The number of potentially novel effectors in *B. lactucae* and *P. infestans* that were identified is much larger than previous reports of effector repertoires from these two species (Fletcher et al. 2019; Haas et al. 2009). These predicted effector repertoires are larger because EffectorO-ML uses N-terminal biochemical characteristics rather than specific amino acid motifs and thus finds effector candidates that would have been missed by prediction pipelines that rely on specific sequences. As with any prediction, some of the candidates will be false positives; therefore, functional characterization is necessary to confirm that they are effectors. The large size of these sets of effector candidates should not preclude functional analysis when combined with additional data from the pathogen of interest such as avirulence gene mapping, population genomics, and gene expression data.

EffectorO-ML was trained on the N-terminal sequences of validated effectors compared to conserved secreted proteins (presumed non-effectors), which, as we show, differ in several biochemical characteristics. Sequences in the N-terminus of effector proteins are known to be important for secretion and translocation of effectors; here we demonstrate that they have distinct biochemical signatures that can be used to predict the probability that a given sequence is an effector. Our positive training set showed increased N-terminal intrinsic disorder compared to conserved proteins, which has also been observed for oomycete RXLR effectors (Marin et al. 2013; Shen et al. 2017). Intrinsically disordered regions are flexible segments of proteins that may adopt changeable conformations upon binding to interacting partners or in changing conditions, such as pH (Marin et al. 2013). It has also been observed that the N-terminus of cytoplasmic effectors are enriched for positively charged amino acids (Sperschneider and Dodds 2022). These observations are intriguing and should be followed up by mutational studies *in vitro* and *in vivo* to clarify their role in effector translocation and/or function.

EffectorO-ML was able to recover the majority of predicted RXLR-EER and WY domain-containing proteins from *B. lactucae* and *P. infestans* without relying on using those motifs for prediction. By focusing on the N-terminus for effector classification, our pipeline allows for a diversity of C-terminal domains, which are typically the functional region of effector proteins. Because EffectorO does not rely on known motifs, it may help to uncover new effector families. Interestingly, the majority of EffectorO-ML candidates had no functional domains predicted using Pfam. This does not mean that the effectors do not contain domains with biochemical activities, but rather their domains have yet to be characterized. Functional validation followed by structural characterization using cryo-EM or x-ray crystallography can be done on these predicted effector proteins, as has been done for WY-containing effectors (Boutemy et al. 2011). The predicted effector repertoires should also be analyzed with the latest machine learning structural prediction algorithms such as trRosetta (Yang et al. 2020), AlphaFold (Jumper et al. 2021), or RoseTTAfold (Baek et al. 2021). Recently, trRosetta was used to predict effectors based on structure for the fungal species *Magnaporthe oryzae* (Seong and Krasileva 2021) and a similar analysis should be done with EffectorO candidates.

The most accurate model for EffectorO-ML was the Random Forest classifier. Random Forest classifiers work by building many decision trees and then taking the consensus of all of the decision trees to calculate an overall probability for a given classification (Pavlov 2000). Because the final output is the average of many independent models, Random Forest classifiers are robust to individual errors in any given single model. The fungal and oomycete effector classifier, EffectorP 3.0, uses a similar approach by utilizing an ensemble classifier that takes the consensus predictions of multiple Naive-Bayes and decision trees to obtain a balanced accuracy of 85% for predicting apoplastic and cytoplasmic effectors. While EffectorP is trained on a larger set of effectors than EffectorO and relies on additional biochemical characteristics (15 features vs. our six), we achieved a similar accuracy to EffectorP 3.0 in predicting oomycete effectors. Our Random Forest model had an estimated 84% accuracy and 82% specificity at a probability cutoff of 0.5. The new EffectorP 3.0 classifies 86/92 (93.5%) of the oomycete effectors used in our positive training set as effectors and 19/88 (21.6%) of our negative training set as effectors, giving it similar sensitivity and specificity to our pipeline. However, predictions did not completely overlap between the two methods; therefore, the two methods could be used as complementary tools to predict high confidence effectors. For example, EffectorP 3.0 does not predict BLE01 from B. lactucae to be an effector, however in this paper we show that it causes cell death in lettuce and is an Avr candidate. EffectorP 3.0 also predicts many more effectors compared to EffectorO, especially in downy mildew genomes (with 40–50% of the secretome predicted as effectors), which could be indicative of a higher number of false positives than EffectorO-ML.

The false positive rate for EffectorO-ML can be decreased by increasing the probability cutoff for classification of an effector; however, this will also result in more false negatives (missed true effectors). For inclusive searches (especially if combined with other lines of evidence), we recommend 0.5 as the threshold for analyses of oomycete genomes. For a slightly less inclusive but more accurate threshold, we recommend a threshold of 0.6. The choice of threshold cutoff depends on downstream effector prediction steps and throughput for effector validation. It may also be possible to further improve the accuracy and specificity of EffectorO by adding additional proteins to the training set or by adding more biochemical and structural characteristics. Gene models may be used instead of ORFs to reduce false positives, although gene prediction algorithms may miss true effectors (as was the case for the *Avr6* candidate *BLE01*). We also used predicted secreted proteins as our input (predicted using the most sensitive version of SignalP 4.1); however, since some effectors may have non-conventional secretion signals, total proteomes could also be used for EffectorO-ML, although there may be an increased false positive rate using this approach and candidates should be evaluated using additional lines of evidence to find true effectors.

Similar to EffectorO-ML, EffectorO-LSP will predict proteins other than effectors, not only due to false positives but also due to the biological reality that there may be lineage-specific proteins that have other functions. Thus, EffectorO-LSP predictions are most useful in combination with other lines of evidence of effector activity (such as EffectorO-ML, expression during infection, RXLR motifs and/or WY domains). It is important to note that CRN effectors do not appear to be lineage-specific (they are conserved across species), so the LSP pipeline may filter out CRN type effectors. Approximately 41% of the *P. infestans* secretome and 34% of the *B. lactucae* secretome were predicted to be lineage-specific. Some of these predicted proteins may not actually be lineage-specific; some may be encoded by pseudogenes or result from failure to detect homology (Weisman et al. 2020). We found that many were expressed in *B. lactucae* during infection, suggesting that some represent true genes. In addition to effectors, these lineage-specific secreted proteins could also be related to the lifestyles of these organisms, their sexual reproduction, or may be products of co-evolution with other microbes rather than co-evolution with the host.

The number of lineage-specific genes detected for a given organism will depend on whether or not closely related organisms are included in the analysis. The user can decide which closely related species to include in order to adjust the stringency of the filter. We recommend that BLAST results from very closely related species not be included in order to reduce the false negative rate. The researcher can choose their own trade-off between false negatives and false positives. If closely related species are included in the analysis, then some true effectors will likely be filtered out due to the presence of effector orthologs in closely related genomes. On the other hand, when only distantly related species are present in the database, there may be more false positives. Note that for species that do not have complete and well annotated genomes compared to closely related organisms, sequences may be inaccurately classified as “species-specific;” however, they can still be considered “lineage-specific” to the other genomes in the database used for comparison. Therefore, we consider the lack of orthologs in other oomycete genomes as a useful characteristic to help predict effectors, even if its accuracy may be limited by the contemporary genomic resources for a given species.

EffectorO was validated for a predicted effector from *B. lactucae,* which was within the region co-segregating with the *Avr6* phenotype, but did not have RXLR or EER motifs or WY domains. We showed that the gene, *BLE01*, induced cell death when expressed in lettuce cultivars expressing the *Dm6* gene but not on cultivars containing other known *Dm* genes, making it a good candidate for *Avr6.* Expression of *BLE01* also caused cell death on RYZ2164. The background of RYZ2164 is proprietary, so it is unknown whether it has *Dm6* in its lineage or another *Dm* gene that recognizes the effector. Bioinformatic analysis of BLE01 revealed that it was similar in sequence and structure to the WY domain containing protein BSW13. Subsequently, when the bias filter was turned off in HMMer, BLE01 was predicted to be a WY domain protein, albeit a low scoring one. Since the filters in HMMer exist to increase the speed of the search at the expense of false negatives, we recommend that HMMer users who are not concerned with speed turn off the bias filter using the option (--nobias) or all filters using the option (--max).

EffectorO-ML and -LSP can expand the set of predicted effectors for identification of genes for resistance to agriculturally important oomycetes. These predictions can be used to choose candidates for “effectoromics” screens (Vleeshouwers and Oliver 2015) to identify effectors with avirulence and virulence activities. For pathogen species without as extensive genetic resources as *B. lactucae*, other lines of evidence can be used to narrow down the list of effectors for functional characterization, including transcriptomic analyses to identify proteins expressed during infection and population genomics to identify loci under selection. Effectors encoded by avirulence genes are valuable for discovering, mapping, and characterizing host resistance genes. Knowledge of R gene–effector interactions will enable the stacking and maintenance of multiple R genes during introgression from wild relatives and can help prevent redundancy of effector recognition specificities within a breeding program. In addition, more accurate annotation of pathogen genomes will enable pathogen-informed R gene deployment and sequence-based monitoring of pathogen populations (Michelmore et al. 2013).

## Acknowledgements

The authors would like to thank John Emerson for greenhouse support, Elizabeth Georgian for manuscript editing and formatting, Kyle Fletcher for very helpful comments and suggestions, and Alexi Balmuth and Mary Corrigan for additional editing. This work was supported by a USDA-AFRI Predoctoral Fellowship to Kelsey Wood (grant no. 2018-67011-28053/proposal no. <u>2017-07123</u>) from the USDA National Institute of Food and Agriculture.

## Supplementary Files

**Supplemental Table 1:** Validated oomycete effector sequences used for the EffectorO-ML positive training set

**Supplemental Table 2:** Information on and citations for the 28 oomycete genomes used in this paper

**Supplemental Table 3:** Protein family domain predictions for the B. lactucae EffectorO-ML predictions

## Notes

https://github.com/mjnur/oomycete-effector-prediction

### Competing Interest Statement

The authors have declared no competing interest.

### Summary of Updates

We have added an analysis of predicted protein domains for EffectorO candidates and have added a comparison with EffectorP 3.0. We also added analysis of CRN type effectors. Minor edits have also been made to the text.

## Literature Cited

Amaro, T. M. M. M., Thilliez, G. J. A., Motion, G. B., and Huitema, E. 2017. A perspective on CRN proteins in the genomics age: Evolution, classification, delivery and function revisited. Front. Plant Sci. 8:1–12

Baek, M., DiMaio, F., Anishchenko, I., Dauparas, J., Ovchinnikov, S., Lee, G. R., Wang, J., Cong, Q., Kinch, L. N., Dustin Schaeffer, R., Millán, C., Park, H., Adams, C., Glassman, C. R., DeGiovanni, A., Pereira, J. H., Rodrigues, A. V., Van Dijk, A. A., Ebrecht, A. C., Opperman, D. J., Sagmeister, T., Buhlheller, C., Pavkov-Keller, T., Rathinaswamy, M. K., Dalwadi, U., Yip, C. K., Burke, J. E., Christopher Garcia, K., Grishin, N. V., Adams, P. D., Read, R. J., and Baker, D. 2021. Accurate prediction of protein structures and interactions using a three-track neural network. Science (80-.). 373:871–876

Bailey, K., Cevik, V., Holton, N., Byrne-Richardson, J., Sohn, K. H., Coates, M., Woods-Tör, A., Aksoy, H. M., Hughes, L., Baxter, L., Jones, J. D. G., Beynon, J., Holub, E. B., and Tör, M. 2011. Molecular cloning of ATR5(Emoy2) from *Hyaloperonospora arabidopsidis*, an avirulence determinant that triggers RPP5-mediated defense in Arabidopsis. Molecular Plant-Microbe Interactions. 24:827–838

Baldauf, S. L. 2003. The Deep Roots of Eukaryotes. Science. 300:1703–1706

Balmer, E. A., and Faso, C. 2021. The Road Less Traveled? Unconventional Protein Secretion at Parasite–Host Interfaces. Front. Cell Dev. Biol. 9:1–12

Baxter, L., Tripathy, S., Ishaque, N., Boot, N., Cabral, A., Kemen, E., Thines, M., Ah-Fong, A., Anderson, R., Badejoko, W., Bittner-Eddy, P., Boore, J. L., Chibucos, M. C., Coates, M., Dehal, P., Delehaunty, K., Dong, S., Downton, P., Dumas, B., Fabro, G., Fronick, C., Fuerstenberg, S. I., Fulton, L., Gaulin, E., Govers, F., Hughes, L., Humphray, S., Jiang, R. H. Y., Judelson, H., Kamoun, S., Kyung, K., Meijer, H., Minx, P., Morris, P., Nelson, J., Phuntumart, V., Qutob, D., Rehmany, A., Rougon-Cardoso, A., Ryden, P., Torto-Alalibo, T., Studholme, D., Wang, Y., Win, J., Wood, J., Clifton, S. W., Rogers, J., Van den Ackerveken, G., Jones, J. D. G., McDowell, J. M., Beynon, J., and Tyler, B. M. 2010. Signatures of adaptation to obligate biotrophy in the *Hyaloperonospora arabidopsidis* genome. Science. 330:1549–1551

Birch, P. R. J., Armstrong, M., Bos, J., Boevink, P., Gilroy, E. M., Taylor, R. M., Wawra, S., Pritchard, L., Conti, L., Ewan, R., Whisson, S. C., van West, P., Sadanandom, A., and Kamoun, S. 2009. Towards understanding the virulence functions of RXLR effectors of the oomycete plant pathogen *Phytophthora infestans*. Journal of Experimental Botany. 60:1133–1140

Bos, J. I. B., Armstrong, M. R., Gilroy, E. M., Boevink, P. C., Hein, I., Taylor, R. M., Zhendong, T., Engelhardt, S., Vetukuri, R. R., Harrower, B., Dixelius, C., Bryan, G., Sadanandom, A., Whisson, S. C., Kamoun, S., and Birch, P. R. J. 2010. *Phytophthora infestans* effector AVR3a is essential for virulence and manipulates plant immunity by stabilizing host E3 ligase CMPG1. Proceedings of the National Academy of Sciences. 107:9909–9914

Boutemy, L. S., King, S. R. F., Win, J., Hughes, R. K., Clarke, T. A., Blumenschein, T. M. A., Kamoun, S., and Banfield, M. J. 2011. Structures of *Phytophthora* RXLR effector proteins: a conserved but adaptable fold underpins functional diversity. Journal of Biological Chemistry. 286:35834–35842

Chou, S., Krasileva, K. V, Holton, J. M., Steinbrenner, A. D., Alber, T., and Staskawicz, B. J. 2011. *Hyaloperonospora arabidopsidis ATR1* effector is a repeat protein with distributed recognition surfaces. Proceedings of the National Academy of Sciences. 108:13323–13328

Deb, D., Anderson, R. G., How-Yew-Kin, T., Tyler, B. M., and McDowell, J. M. 2018. Conserved RxLR Effectors From Oomycetes *Hyaloperonospora arabidopsidis* and *Phytophthora sojae* Suppress PAMP- and Effector-Triggered Immunity in Diverse Plants. Molecular Plant-Microbe Interactions. 31:374–385

Derevnina, L., Petre, B., Kellner, R., Dagdas, Y. F., Sarowar, M. N., Giannakopoulou, A., De la Concepcion, J. C., Chaparro-Garcia, A., Pennington, H. G., van West, P., and Kamoun, S. 2016. Emerging oomycete threats to plants and animals. Philos. Trans. R. Soc. Lond. B Biol. Sci. 371

Dong, S., Raffaele, S., and Kamoun, S. 2015. The two-speed genomes of filamentous pathogens: waltz with plants. Current Opinion in Genetics & Development. 35:57–65

Dodds, P. N., Lawrence, G. J., Catanzariti, A. M., Teh, T., Wang, C. I. A., Ayliffe, M. A., Kobe, B., and Ellis, J. G. 2006. Direct protein interaction underlies gene-for-gene specificity and coevolution of the flax resistance genes and flax rust avirulence genes. Proceedings of the National Academy of Sciences. 103:8888–8893

Dou, D., Kale, S. D., Wang, X., Jiang, R. H. Y., Bruce, N. A., Arredondo, F. D., Zhang, X., and Tyler, B. M. 2008. RXLR-mediated entry of *Phytophthora sojae* effector Avr1b into soybean cells does not require pathogen-encoded machinery. Plant Cell. 20:1930– 1947

Dunker, A.K., Lawson, J.D., Brown, C.J., Williams, R.M., Romero, P., Oh, J.S., Oldfield, C.J., Campen, A.M., Ratliff, C.M., Hipps, K.W., Ausio, J., Nissen, M.S., Reeves, R., Kang, C.H., Kissinger, C.R., Bailey, R.W., Griswold, M.D., Chiu, W., Garner, E.C. and Obradovic, Z. 2001. Intrinsically disordered protein. Journal of Molecular Graphics and Modelling. 19, 26–59

Eddy, S. R. 2011. Accelerated Profile HMM Searches. PLoS Comput. Biol. 7:e1002195

Ellis, J. G., and Dodds, P. N. 2011. Showdown at the RXLR motif: Serious differences of opinion in how effector proteins from filamentous eukaryotic pathogens enter plant cells. Proceedings of the National Academy of Sciences. 108:14381–14382

ExPaSy (Expert Protein Analysis System). 2004. Encyclopedic Dictionary of Genetics, Genomics and Proteomics.

Fabro, G., Steinbrenner, J., Coates, M., Ishaque, N., Baxter, L., Studholme, D. J., Körner, E., Allen, R. L., Piquerez, S. J. M., Rougon-Cardoso, A., Greenshields, D., Lei, R., Badel, J. L., Caillaud, M.-C., Sohn, K.-H., Van den Ackerveken, G., Parker, J. E., Beynon, J., and Jones, J. D. G. 2011. Multiple Candidate Effectors from the Oomycete Pathogen Hyaloperonospora arabidopsidis Suppress Host Plant Immunity. PLOS Pathogens. 7:e1002348

Fauchere, J.L. and Pliska, V.E. 1983. Hydrophobic parameters pi of amino acid side chains from partitioning of N-acetyl-amino-acid amides. European Journal of Medicinal Chemistry. 18, 369–375

Fletcher, K., Gil, J., Bertier, L. D., Kenefick, A., Wood, K. J., Zhang, L., Reyes-Chin-Wo, S., Cavanaugh, K., Tsuchida, C., Wong, J., and Michelmore, R. 2019. Genomic signatures of heterokaryosis in the oomycete pathogen *Bremia lactucae*. Nature Communications. 10:2645

Fu, L., Niu, B., Zhu, Z., Wu, S., and Li, W. 2012. CD-HIT: accelerated for clustering the next-generation sequencing data. Bioinformatics. 28:3150–3152

Giesbers, A. K. J., Pelgrom, A. J. E., Visser, R. G. F., Niks, R. E., Van den Ackerveken, G., and Jeuken, M. J. W. 2017. Effector-mediated discovery of a novel resistance gene against *Bremia lactucae* in a nonhost lettuce species. New Phytologist. 216:915–926

Godfrey, D., Böhlenius, H., Pedersen, C., Zhang, Z., Emmersen, J., and Thordal-Christensen, H. 2010. Powdery mildew fungal effector candidates share N-terminal Y/F/WxC-motif. BMC Genomics. 11:317

Gu, B., Kale, S. D., Wang, Q., Wang, D., Pan, Q., Cao, H., Meng, Y., Kang, Z., Tyler, B. M., and Shan, W. 2011. Rust Secreted Protein Ps87 Is Conserved in Diverse Fungal Pathogens and Contains a RXLR-like Motif Sufficient for Translocation into Plant Cells. PLOS ONE. 6:e27217

Haas, B. J., Kamoun, S., Zody, M. C., Jiang, R. H. Y., Handsaker, R. E., Cano, L. M., Grabherr, M., Kodira, C. D., Raffaele, S., Torto-Alalibo, T., Bozkurt, T. O., Ah-Fong, A. M. V., Alvarado, L., Anderson, V. L., Armstrong, M. R., Avrova, A., Baxter, L., Beynon, J., Boevink, P. C., Bollmann, S. R., Bos, J. I. B., Bulone, V., Cai, G., Cakir, C., Carrington, J. C., Chawner, M., Conti, L., Costanzo, S., Ewan, R., Fahlgren, N., Fischbach, M. A., Fugelstad, J., Gilroy, E. M., Gnerre, S., Green, P. J., Grenville-Briggs, L. J., Griffith, J., Grünwald, N. J., Horn, K., Horner, N. R., Hu, C.-H., Huitema, E., Jeong, D.-H., Jones, A. M. E., Jones, J. D. G., Jones, R. W., Karlsson, E. K., Kunjeti, S. G., Lamour, K., Liu, Z., Ma, L., Maclean, D., Chibucos, M. C., McDonald, H., McWalters, J., Meijer, H. J. G., Morgan, W., Morris, P. F., Munro, C. A., O’Neill, K., Ospina-Giraldo, M., Pinzón, A., Pritchard, L., Ramsahoye, B., Ren, Q., Restrepo, S., Roy, S., Sadanandom, A., Savidor, A., Schornack, S., Schwartz, D. C., Schumann, U. D., Schwessinger, B., Seyer, L., Sharpe, T., Silvar, C., Song, J., Studholme, D. J., Sykes, S., Thines, M., van de Vondervoort, P. J. I., Phuntumart, V., Wawra, S., Weide, R., Win, J., Young, C., Zhou, S., Fry, W., Meyers, B. C., van West, P., Ristaino, J., Govers, F., Birch, P. R. J., Whisson, S. C., Judelson, H. S., and Nusbaum, C. 2009. Genome sequence and analysis of the Irish potato famine pathogen *Phytophthora infestans*. Nature. 461:393–398

Haverkort, A. J., Boonekamp, P. M., Hutten, R., Jacobsen, E., Lotz, L. A. P., Kessel, G. J. T., Visser, R. G. F., and van der Vossen, E. A. G. 2008. Societal Costs of Late Blight in Potato and Prospects of Durable Resistance Through Cisgenic Modification. Potato Research. 51:47–57

He, J., Ye, W., Choi, D. S., Wu, B., Zhai, Y., Guo, B., Duan, S., Wang, Y., Gan, J., Ma, W., and Ma, J. 2019. Structural analysis of *Phytophthora* suppressor of RNA silencing 2 (PSR2) reveals a conserved modular fold contributing to virulence. Proceedings of the National Academy of Sciences. 116:8054–8059

Janin, J. 1979. Surface and inside volumes in globular proteins. Nature. 277, 491– 492

Jiang, R. H. Y., de Bruijn, I., Haas, B. J., Belmonte, R., Löbach, L., Christie, J., van den Ackerveken, G., Bottin, A., Bulone, V., Díaz-Moreno, S. M., Dumas, B., Fan, L., Gaulin, E., Govers, F., Grenville-Briggs, L. J., Horner, N. R., Levin, J. Z., Mammella, M., Meijer, H. J. G., Morris, P., Nusbaum, C., Oome, S., Phillips, A. J., van Rooyen, D., Rzeszutek, E., Saraiva, M., Secombes, C. J., Seidl, M. F., Snel, B., Stassen, J. H. M., Sykes, S., Tripathy, S., van den Berg, H., Vega-Arreguin, J. C., Wawra, S., Young, S. K., Zeng, Q., Dieguez-Uribeondo, J., Russ, C., Tyler, B. M., and van West, P. 2013. Distinctive expansion of potential virulence genes in the genome of the oomycete fish pathogen *Saprolegnia parasitica*. PLOS Genetics. 9:e1003272

Jones, S. and Thornton, J.M. 1997. Analysis of protein-protein interaction sites using surface patches. Journal of Molecular Biology. 272, 121–132

Jumper, J., Evans, R., Pritzel, A., Green, T., Figurnov, M., Ronneberger, O., Tunyasuvunakool, K., Bates, R., Žídek, A., Potapenko, A., Bridgland, A., Meyer, C., Kohl, S. A. A., Ballard, A. J., Cowie, A., Romera-Paredes, B., Nikolov, S., Jain, R., Adler, J., Back, T., Petersen, S., Reiman, D., Clancy, E., Zielinski, M., Steinegger, M., Pacholska, M., Berghammer, T., Bodenstein, S., Silver, D., Vinyals, O., Senior, A. W., Kavukcuoglu, K., Kohli, P., and Hassabis, D. 2021. Highly accurate protein structure prediction with AlphaFold. Nature. 596:583–589

Kale, S. D., Gu, B., Capelluto, D. G. S., Dou, D., Feldman, E., Rumore, A., Arredondo, F. D., Hanlon, R., Fudal, I., Rouxel, T., Lawrence, C. B., Shan, W., and Tyler, B. M. 2010. External Lipid PI3P Mediates Entry of Eukaryotic Pathogen Effectors into Plant and Animal Host Cells. Cell. 142:284–295

King, S. R. F., McLellan, H., Boevink, P. C., Armstrong, M. R., Bukharova, T., Sukarta, O., Win, J., Kamoun, S., Birch, P. R. J., and Banfield, M. J. 2014. *Phytophthora infestans* RXLR effector PexRD2 interacts with host MAPKKK ε to suppress plant immune signaling. Plant Cell. 26:1345–1359

Kyte, J. and Doolittle, R.F. 1982. A simple method for displaying the hydropathic character of a protein. Journal of Molecular. Biology. 157, 105–132

Leonelli, L., Pelton, J., Schoeffler, A., Dahlbeck, D., Berger, J., Wemmer, D. E., and Staskawicz, B. 2011. Structural elucidation and functional characterization of the Hyaloperonospora arabidopsidis effector protein ATR13. PLoS Pathogens. 7:e1002428

Li, H., Handsaker, B., Wysoker, A., Fennell, T., Ruan, J., Homer, N., Marth, G., Abecasis, G., and Durbin, R. 2009. The Sequence Alignment/Map format and SAMtools. Bioinformatics. 25:2078–2079

Li, W., and Godzik, A. 2006. Cd-hit: a fast program for clustering and comparing large sets of protein or nucleotide sequences. Bioinformatics. 22:1658–1659

Liu, T., Song, T., Zhang, X., Yuan, H., Su, L., Li, W., Xu, J., Liu, S., Chen, L., Chen, T., Zhang, M., Gu, L., Zhang, B., and Dou, D. 2014. Unconventionally secreted effectors of two filamentous pathogens target plant salicylate biosynthesis. Nature Communications. 5

Marín, M., Uversky, V. N., and Ott, T. 2013. Intrinsic disorder in pathogen effectors: Protein flexibility as an evolutionary hallmark in a molecular arms race. Plant Cell. 25:3153– 3157

McGowan, J., and Fitzpatrick, D. A. 2017. Genomic, Network, and Phylogenetic Analysis of the Oomycete Effector Arsenal. mSphere. 2

Michelmore, R. W., Christopoulou, M., and Caldwell, K. S. 2013. Impacts of resistance gene genetics, function, and evolution on a durable future. Annual Review of Phytopathology 51:291–319

Mistry, J., Chuguransky, S., Williams, L., Qureshi, M., Salazar, G. A., Sonnhammer, E. L. L., Tosatto, S. C. E., Paladin, L., Raj, S., Richardson, L. J., Finn, R. D., and Bateman, A. 2021. Pfam: The protein families database in 2021. Nucleic Acids Res. 49:D412–D419

Pavlov, Y. L. 2000. Random Forests. VSP Publishers, Utrecht, the Netherlands.

Pecrix, Y., Buendia, L., Penouilh-Suzette, C., Maréchaux, M., Legrand, L., Bouchez, O., Rengel, D., Gouzy, J., Cottret, L., Vear, F., and Godiard, L. 2019. Sunflower resistance to multiple downy mildew pathotypes revealed by recognition of conserved effectors of the oomycete *Plasmopara halstedii*. Plant Journal. 97:730–748

Pedregosa F, Varoquaux G, Gramfort A, Michel V, Thirion B, Grisel O, Blondel, M., Prettenhofer, P., Weiss, R., Dubourg, V., Vanderplas, J., Passos, A., Cournapeau, D., Brucher, M., Perrot, M., and Duchesnay, E. Scikit-learn: Machine Learning in Python. Journal of Machine Learning Research 2011;12: 2825–2830.

Pelgrom, A. J. E., Eikelhof, J., Elberse, J., Meisrimler, C.-N., Raedts, R., Klein, J., and Van den Ackerveken, G. 2019. Recognition of lettuce downy mildew effector BLR38 in *Lactuca serriola* LS102 requires two unlinked loci. Molecular Plant Pathology. 20:240–253

Pel, M. A., Foster, S. J., Park, T.-H., Rietman, H., van Arkel, G., Jones, J. D. G., Van Eck, H. J., Jacobsen, E., Visser, R. G. F., and Van der Vossen, E. A. G. 2009. Mapping and cloning of late blight resistance genes from *Solanum venturii* using an interspecific candidate gene approach. Molecular Plant-Microbe Interactions. 22:601–615

Peng, K., Radivojac, P., Vucetic, S., Dunker, A. K., and Obradovic, Z. 2006. Length-dependent prediction of protein intrinsic disorder. BMC Bioinformatics. 7:208

Petersen, T. N., Brunak, S., von Heijne, G., and Nielsen, H. 2011. SignalP 4.0: discriminating signal peptides from transmembrane regions. Nature Methods. 8:785–786

Rehmany, A. P., Gordon, A., Rose, L. E., Allen, R. L., Armstrong, M. R., Whisson, S. C., Kamoun, S., Tyler, B. M., Birch, P. R. J., and Beynon, J. L. 2005. Differential recognition of highly divergent downy mildew avirulence gene alleles by RPP1 resistance genes from two Arabidopsis lines. Plant Cell. 17:1839–1850

Rujirawat, T., Patumcharoenpol, P., Kittichotirat, W., and Krajaejun, T. 2019. Oomycete Gene Table: an online database for comparative genomic analyses of the oomycete microorganisms. Database. 2019

Sayers, E. W., Beck, J., Brister, J. R., Bolton, E. E., Canese, K., Comeau, D. C., Funk, K., Ketter, A., Kim, S., Kimchi, A., Kitts, P. A., Kuznetsov, A., Lathrop, S., Lu, Z., McGarvey, K., Madden, T. L., Murphy, T. D., O’Leary, N., Phan, L., Schneider, V. A., Thibaud-Nissen, F., Trawick, B. W., Pruitt, K. D., and Ostell, J. 2020. Database resources of the National Center for Biotechnology Information. Nucleic Acids Research. 48:D9–D16

Seong, K., and Krasileva, K. V. 2021. Computational structural genomics unravels common folds and predicted functions in the secretome of fungal phytopathogen Magnaporthe oryzae. bioRxiv doi: 10.1101/2021.01.25.427855

Sharma, R., Xia, X., Cano, L. M., Evangelisti, E., Kemen, E., Judelson, H., Oome, S., Sambles, C., van den Hoogen, D. J., Kitner, M., Klein, J., Meijer, H. J. G., Spring, O., Win, J., Zipper, R., Bode, H. B., Govers, F., Kamoun, S., Schornack, S., Studholme, D. J., Van den Ackerveken, G., and Thines, M. 2015. Genome analyses of the sunflower pathogen *Plasmopara halstedii* provide insights into effector evolution in downy mildews and *Phytophthora*. BMC Genomics. 16:741

Shen, D., Li, Q., Sun, P., Zhang, M., and Dou, D. 2017. Intrinsic disorder is a common structural characteristic of RxLR effectors in oomycete pathogens. Fungal Biology. 121:911–919

Sperschneider, J., and Dodds, P. N. 2022. EffectorP 3.0: Prediction of Apoplastic and Cytoplasmic Effectors in Fungi and Oomycetes. Molecular Plant-Microbe Interactions. 35:146–156

Sperschneider, J., Dodds, P. N., Gardiner, D. M., Singh, K. B., and Taylor, J. M. 2018. Improved prediction of fungal effector proteins from secretomes with EffectorP 2.0. Molecular Plant Pathology. 19:2094–2110

Sperschneider, J., Gardiner, D. M., Dodds, P. N., Tini, F., Covarelli, L., Singh, K. B., Manners, J. M., and Taylor, J. M. 2016. EffectorP: predicting fungal effector proteins from secretomes using machine learning. New Phytologist. 210:743–761

Stajich JE, Block D, Boulez K, Brenner SE, Chervitz SA, Dagdigian C, Fuellen G, Gilbert JG, Korf I, Lapp H, Lehväslaiho H, Matsalla C, Mungall CJ, Osborne BI, Pocock MR, Schattner P, Senger M, Stein LD, Stupka E, Wilkinson MD, and Birney E. The Bioperl toolkit: Perl modules for the life sciences. Genome Res. 2002 Oct;12(10):1611–8.

Stassen, J. H. M., den Boer, E., Vergeer, P. W. J., Andel, A., Ellendorff, U., Pelgrom, K., Pel, M., Schut, J., Zonneveld, O., Jeuken, M. J. W., and Van den Ackerveken, G. 2013. Specific In Planta Recognition of Two GKLR Proteins of the Downy Mildew *Bremia lactucae* Revealed in a Large Effector Screen in Lettuce. Molecular Plant-Microbe Interactions. 26:1259–1270

Thompson, J. D., Higgins, D. G., and Gibson, T. J. 1994. CLUSTAL W: Improving the sensitivity of progressive multiple sequence alignment through sequence weighting, position-specific gap penalties and weight matrix choice. Nucleic Acids Research. 22:4673– 4680

Urban, M., Cuzick, A., Seager, J., Wood, V., Rutherford, K., Venkatesh, S. Y., De Silva, N., Martinez, M. C., Pedro, H., Yates, A. D., Hassani-Pak, K., and Hammond-Kosack, K. E. 2019. PHI-base: the pathogen–host interactions database. Nucleic Acids Research.

Vleeshouwers, V. G. A. A., and Oliver, R. P. 2015. Effectors as Tools in Disease Resistance Breeding Against Biotrophic, Hemibiotrophic, and Necrotrophic Plant Pathogens. Molecular Plant-Microbe Interactions. 2015:17–27

Wang, Y., Tyler, B. M., and Wang, Y. 2019. Defense and Counterdefense During Plant-Pathogenic Oomycete Infection. Annual. Review of Microbiology. 73:667–696

Weisman, C. M., Murray, A. W., and Eddy, S. R. 2020. Many but not all lineage-specific genes can be explained by homology detection failure. PLOS Biology 18(11):e3000862.

Whisson, S. C., Boevink, P. C., Moleleki, L., Avrova, A. O., Morales, J. G., Gilroy, E. M., Armstrong, M. R., Grouffaud, S., van West, P., Chapman, S., Hein, I., Toth, I. K., Pritchard, L., and Birch, P. R. J. 2007. A translocation signal for delivery of oomycete effector proteins into host plant cells. Nature. 450:115–118

Win, J., Morgan, W., Bos, J., Krasileva, K. V, Cano, L. M., Chaparro-Garcia, A., Ammar, R., Staskawicz, B. J., and Kamoun, S. 2007. Adaptive evolution has targeted the C-terminal domain of the RXLR effectors of plant pathogenic oomycetes. Plant Cell. 19:2349–69

Win, J., Krasileva, K. V., Kamoun, S., Shirasu, K., Staskawicz, B. J., and Banfield, M. J. 2012. Sequence Divergent RXLR Effectors Share a Structural Fold Conserved across Plant Pathogenic Oomycete Species. PLOS Pathogens. 8:e1002400

Wood, K., Nur, M., Gil, J., Fletcher, K., Lakeman, K., Gothberg, A., Khuu, T., Kopetzky, J., Pandya, A., Pel, M., and Michelmore, R. 2020. Effector prediction and characterization in the oomycete pathogen *Bremia lactucae* reveal host-recognized WY domain proteins that lack the canonical RXLR motif. PLOS Pathogens 10: e1009012

Wroblewski T., Caldwell K .S., Piskurewicz U., Cavanaugh K. A., Xu H., Kozik A., Ochoa O., McHale L. K., Lahre K., Jelenska J., Castillo J. A., Blumenthal D., Vinatzer B. A., Greenberg J. T., and Michelmore R. W. 2009. Comparative large-scale analysis of interactions between several crop species and the effector repertoires from multiple pathovars of *Pseudomonas* and *Ralstonia*. Plant Physiology. 150: 1733–49. doi:10.1104/pp.109.140251

Yang, J., Anishchenko, I., Park, H., Peng, Z., Ovchinnikov, S., and Baker, D. 2020. Improved protein structure prediction using predicted interresidue orientations. Proceedings of the National Academy of Sciences. 117:1496–1503

Zheng, X., McLellan, H., Fraiture, M., Liu, X., Boevink, P. C., Gilroy, E. M., Chen, Y., Kandel, K., Sessa, G., Birch, P. R. J., and Brunner, F. 2014. Functionally Redundant RXLR Effectors from Phytophthora infestans Act at Different Steps to Suppress Early flg22-Triggered Immunity. PLOS Pathogens. 10:e1004057

Zhou, K., Huang, B., Zou, M., Lu, D., He, S., and Wang, G. 2015. Data in support of genome-wide identification of lineage-specific genes within *Caenorhabditis elegans*. Data in Brief. 4:595–601

Zimmerman, J.M., Eliezer, N. and Simha, R. 1968. The characterization of amino acid sequences in proteins by statistical methods. Journal of Theoretical Biology. 21, 170–201

